# Selection against admixture and gene regulatory divergence in a long-term primate field study

**DOI:** 10.1101/2021.08.19.456711

**Authors:** Tauras P. Vilgalys, Arielle S. Fogel, Raphael S. Mututua, J. Kinyua Warutere, Long’ida Siodi, Jordan A. Anderson, Sang Yoon Kim, Tawni N. Voyles, Jacqueline A. Robinson, Jeffrey D. Wall, Elizabeth A. Archie, Susan C. Alberts, Jenny Tung

**Affiliations:** Department of Evolutionary Anthropology, Duke University, Durham, NC, USA; Section of Genetic Medicine, University of Chicago, Chicago, IL, USA; University Program in Genetics and Genomics, Duke University, Durham, NC, USA; Amboseli Baboon Research Project, Amboseli National Park, Kenya; Institute for Human Genetics, University of California, San Francisco, CA, USA; Department of Biological Sciences, University of Notre Dame, Notre Dame, IN, USA; Department of Biology, Duke University, Durham, NC, USA; Duke University Population Research Institute, Duke University, Durham, NC, USA

**Author notes:** Contributed equally to this work (Tauras P. Vilgalys, Arielle S. Fogel).

## Abstract

Admixture has profoundly influenced evolution across the tree of life, including in humans and other primates^1,2^. However, we have limited insight into the genetic and phenotypic consequences of admixture in primates, especially during its key early stages. Here, we address this gap by combining 50 years of field observations with population and functional genomic data from yellow (*Papio cynocephalus*) and anubis (*P. anubis*) baboons in Kenya, in a longitudinally studied population that has experienced both historical and recent admixture^3^. We use whole-genome sequencing to characterize the extent of the hybrid zone, estimate local ancestry for 442 known individuals, and predict the landscape of introgression across the genome. Despite no major fitness costs to hybrids, we identify signatures of selection against introgression that are strikingly similar to those described for archaic hominins^4–6^. These signatures are strongest near loci with large ancestry effects on gene expression, supporting the importance of gene regulation in primate evolution and the idea that selection targeted large regulatory effects following archaic hominin admixture^7,8^. Our results show that genomic data and field observations of hybrids are important and mutually informative. They therefore demonstrate the value of other primates as living models for phenomena that we cannot observe in our own lineage.

## Main

The past decade of anthropological genomics has revolutionized our understanding of human evolution. It is now clear that the ancestors of modern humans not only intermixed with Neanderthals and other archaic hominins, but that the legacy of this contact continues to shape trait variation in humans today^7,9–13^. Even as these findings reshape our conception of human origins, they also bring us more closely in line with other animals, including our primate relatives, where hybridization is commonly observed in the wild^1^. Studies of other living primates therefore provide valuable context for understanding admixture dynamics in our own lineage. For instance, genomic analyses make clear that selection removed introgressed archaic ancestry from modern human genomes, probably with greatest effect shortly after contact^14–17^. However, we do not know whether these early generation hybrids experienced major fitness costs; whether mate choice, intrinsic incompatibilities, or ecological selection were involved; or whether regulatory variation played an outsized role. These questions can be addressed in nonhuman primate hybrid zones where complementary demographic, phenotypic, and genomic data can be collected.

Studies of primate hybrids in the wild suggest that ancestry frequently predicts variation in morphological, behavioral, and life-history traits, but often does not result in overt fitness costs^18–21^. However, no study to date has coupled field observations with population and functional genomic analysis to investigate the consequences of admixture at both organismal and molecular levels. Such information is crucial for understanding admixture dynamics in primate evolution, including the range of phenotypic outcomes compatible with evidence for selection against introgression in hominins. Long-term field studies of primate hybrid zones also offer the rare opportunity to directly measure the functional consequences of introgressed alleles in wild individuals, and to follow the course of hybridization and natural selection in real-time, across multiple generations. They therefore can provide key insight into the determinants of introgression across the genome and the timescales during which genomic signatures of selection emerge.

Here, we take advantage of such an opportunity by investigating the selective consequences of admixture between yellow baboons (*P. cynocephalus*) and anubis baboons (*P. anubis*)^22^, part of a sub-Saharan radiation of cercopithecine monkeys that has long served as a model for human ecology, evolution, and disease^23^. Anubis and yellow baboons diverged ~1.4 million years ago and occupy large, mostly non-overlapping ranges^24^, but interbreed to produce viable and fertile offspring where their geographic ranges meet^23^ (Fig. 1A). We concentrate on the region in and around the Amboseli basin of southern Kenya, where we have completed fifty years of continuous observation on a population near the center of the hybrid zone^3^. Members of this majority-yellow baboon population include descendants of historical admixture prior to the start of monitoring in 1971, as well as products of a directly observed, recent wave of admixture that began in 1982 (up to 7 baboon generations ago)^25,26^. In Amboseli, hybrids do not experience obvious fitness costs, and anubis ancestry in this population may in fact confer benefits, including accelerated maturation, increased mating success, and an increased rate of opposite sex affiliation, a predictor of longer lifespan in this population^27–30^. However, field data and microsatellite-based analyses indicate that the hybrid zone is geographically constrained^22^. If so, natural selection may also act to limit gene flow between these species—a hypothesis that can be tested at the genomic level.

**Figure 1:**
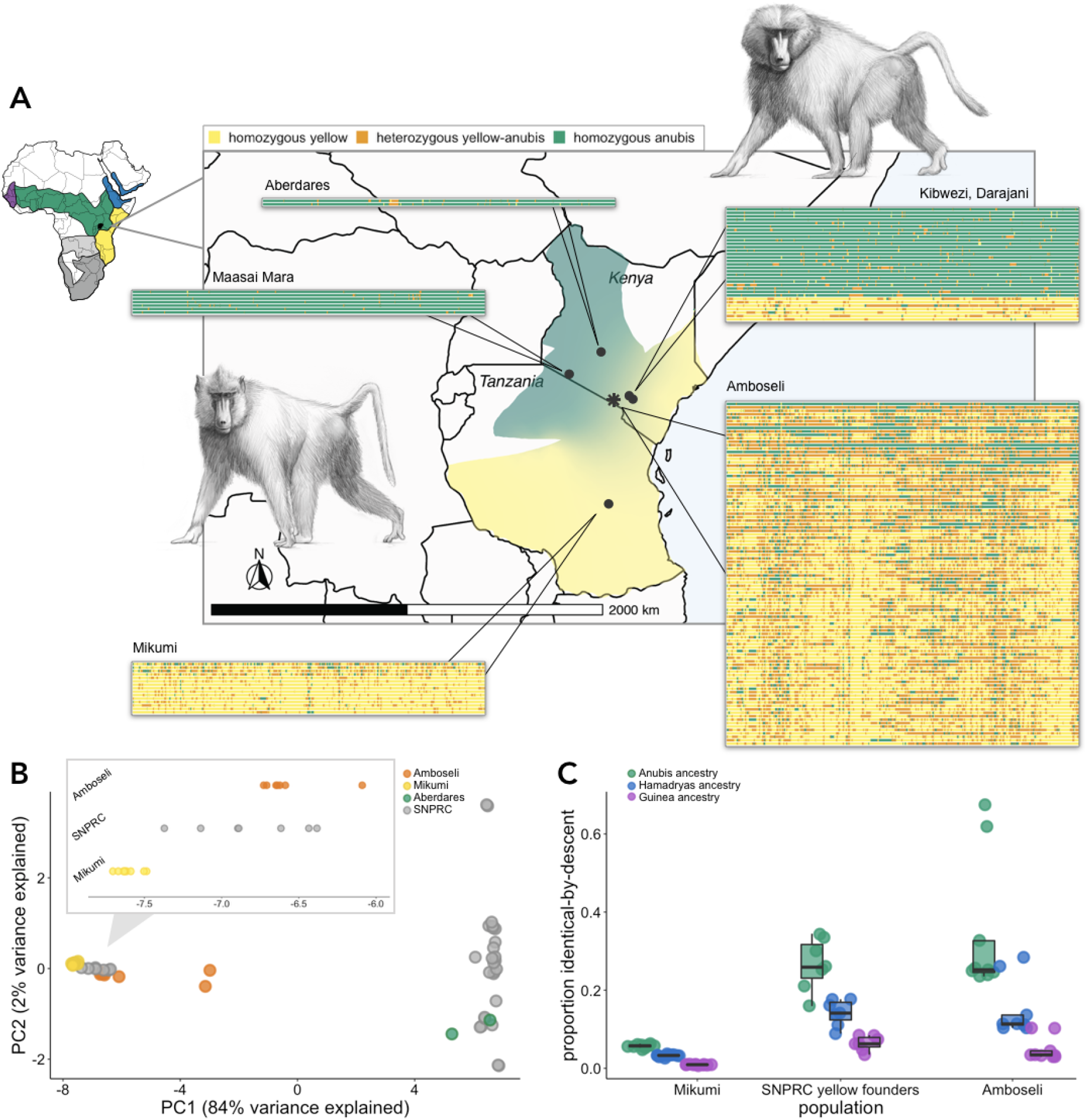
The structure of the baboon hybrid zone in Amboseli and the surrounding region. **(A)** Geographic locations and local genetic ancestry estimates for baboons in this study (Amboseli is denoted by the black asterisk). For each population, each horizontal row corresponds to the first 20 Mb of chromosome 1 for one individual, organized top to bottom from the most anubis to most yellow. For Amboseli, a random subsample of 100 individuals is shown. Background shading shows the ranges of yellow baboons, anubis baboons, and hybrid populations in Kenya and Tanzania (white shading indicates no data or geographic regions baboons do not inhabit)^36^. The ranges of anubis (green), yellow (yellow), hamadryas (blue), Guinea (purple), Kinda (light grey), and chacma (dark grey) in Africa are shown in the upper left-hand map^24^. Baboon drawings by Christopher Smith. **(B)** Principal components (PC) analysis of genotypes from high coverage samples from individuals from SNPRC (grey circles), the Aberdares region (green circles), Mikumi (yellow circles), and Amboseli (orange circles). Inset: close up of “yellow-like” individuals along PC1. SNPRC yellow baboon founders resemble Amboseli yellow baboons and appear to be admixed. **(C)** *IBDMix*^35^ results for three sets of yellow or majority yellow baboons. SNPRC yellow baboon founders exhibit non-negligible identity-by-descent (IBD) with anubis baboons. IBD estimates for SNPRC yellow-anubis are greater than the IBD estimated for two other baboon species that are similarly diverged from yellow baboons, but where gene flow is precluded by geography^24^. The same pattern is observed for majority-yellow baboons in Amboseli, but not for the unadmixed yellow population from Mikumi, Tanzania.

## Structure of the baboon hybrid zone

To assess selection against interbreeding between yellow and anubis baboons, we used whole-genome resequencing data to evaluate genome-wide ancestry patterns for animals sampled in and near the Amboseli hybrid zone (Fig. 1; Table S1). We generated new resequencing data from 430 wild baboons (17 high coverage [mean 22.51x]; 413 low coverage [mean 1.04x]), and combined these data with 36 previously published wild baboon genomes from Amboseli^26,31^ (n=22), the Maasai Mara National Reserve^26^ (n=7), the Aberdares region of central Kenya^24^ (n=2), and Mikumi National Park in Tanzania^24,26^ (n=5). We also included the genomes of 39 captive baboons from the Southwest National Primate Research Center (SNPRC; n=33)^24,32^ and the Washington National Primate Research Center (WNPRC; n=6)^26^, including 31 anubis or yellow founders of the SNPRC colony that were trapped in southern Kenya in the 1960’s^26,33^. We sampled most extensively from the natural hybrid population in Amboseli, where we analyzed 442 individuals born between 1969 (two years before routine observations began) and 2016. Cumulatively, we identified more than 50 million common single nucleotide variants within our sample (minor allele frequency > 5%), substantially adding to the available information on polymorphism and divergence in wild primates.

We estimated global and local ancestry for each individual using a composite likelihood method suitable for low coverage data (*LCLAE*^26^; see Supplementary Methods for simulations and comparison with other methods). Our estimates show that all baboons from Amboseli are admixed (Fig. 1A; mean = 37% genome-wide anubis ancestry ±10% s.d.), including many whose ancestry can be traced through the observational and pedigree data to anubis-like immigrants within the last seven generations. These results are consistent with estimates using *F_4_*-ratios on high-coverage genomes, which produce a mean estimate of 50.8% anubis ancestry for two individuals with recent anubis-like ancestors (versus 51.1% from *LCLAE*) and 23.2% for seven individuals believed to be affected by historical admixture only (versus 24.9% from *LCLAE*) (Supplementary Methods)^34^.

We also confirmed that putatively unadmixed anubis baboon founders of the SNPRC colony, anubis baboons from Maasai Mara, and yellow baboons from Mikumi show little evidence of admixture (Fig. 1, S5). However, our analyses do reveal 10-22% anubis ancestry in the yellow baboons used to found the SNPRC colony, who were previously thought to be unadmixed (Fig. 1A-C)^32^. A reference panel-free approach to identifying introgression (*IBDMix*^35^) confirms this pattern (Fig. 1C). Incomplete lineage sorting is unlikely to explain it, as the degree of identity-by-descent between the yellow baboon SNPRC founders and anubis baboons exceeds the ~10% between the SNPRC yellow founders and other baboon species (Guinea and hamadryas) that share the same divergence time, but whose geographic range makes gene flow with yellow baboons unlikely (Fig. 1A; Supplementary Methods). Combined with evidence for yellow ancestry in an anubis baboon sampled in the Aberdares region in central Kenya, well within the anubis baboon range (Fig. 1A; 15.2% IBD sharing with Mikumi yellow baboons, similar to a previous estimate of 10.5%^24^), our results indicate that gene flow is a common feature of yellow and anubis baboon evolution both within and outside the Amboseli region.

## Selection against introgression in Amboseli

To investigate genomic evidence for selection against yellow-anubis baboon hybridization, we focused on the large, multigenerational data set from Amboseli. We began by replicating three analyses used to infer selection against archaic hominin introgression in humans^4–6^. First, we tested for a relationship between yellow-anubis genetic divergence and anubis ancestry in Amboseli. Since the Amboseli population is largely of yellow baboon origin, if hybridization is deleterious selection is expected to be less permissive of introgressed anubis ancestry with increasing yellow-anubis divergence. Indeed, we find that anubis ancestry is systematically lower in regions of the genome with more fixed differences (Fig. 2A; Spearman’s rho = −0.119, p = 8.05 × 10^−34^; Supplementary Methods), similar to the negative correlation between the density of fixed human-Neanderthal differences and introgressed Neanderthal ancestry in modern humans^5^ (Fig. 2B; Supplementary Methods; Table S2). The mean frequency of anubis baboon alleles in the most diverged percentile of the baboon genome is thus 6.7% lower than the frequency of anubis alleles in the least diverged percentile.

**Figure 2:**
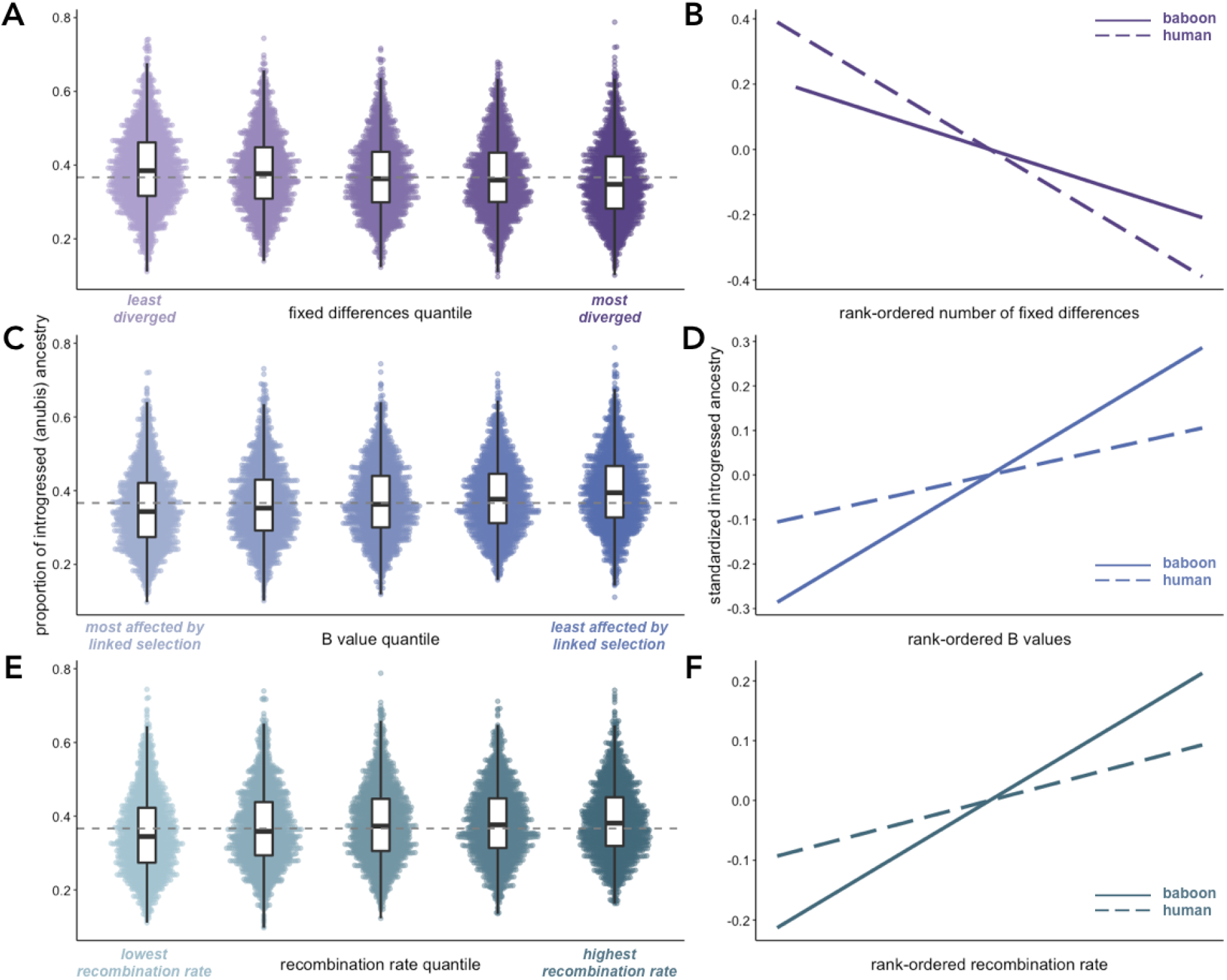
Selection against introgression in the Amboseli baboons mirrors patterns described for archaic hominin admixture. **(A, C, E)** Proportion of introgressed (anubis) ancestry in Amboseli in 250 kb windows (n = 10,324 total windows) as a function of **(A)** fixed differences between yellow and anubis baboons (Spearman’s rho = 0.127, p = 2.49 × 10^−38^), **(C)** mean B statistic value (rho = 0.168, p = 1.73 × 10^−66^), and **(E)** mean recombination rate (rho = −0.119, p = 8.05 × 10^−34^) in each window, divided into quintiles. Dashed grey lines correspond to median anubis ancestry across all windows (0.367). Data are binned into quantiles for visualization purposes only; all analyses were conducted using continuous values. **(B, D, F)** Predicted relationships between introgressed ancestry and all three measures are observed for both anubis ancestry in the Amboseli baboons (solid lines) and Neanderthal ancestry in modern human genomes (dashed lines)^4–6^, consistent with selection against introgression in both cases. Panels show the relationship between introgressed ancestry and **(B)** the rank-ordered number of fixed differences, **(D)** mean B value, and **(F)** mean local recombination rate. Mean introgressed ancestry is centered on 0 and divided by the standard deviation for each species to facilitate comparison.

Second, we tested whether introgressed anubis ancestry is depleted in genomic regions that are likely to be affected by linked selection as summarized by the B statistic^37^ (i.e., due to high gene density per recombination distance). To do so, we recalculated B statistic values for baboons based on the baboon reference genome (Supplementary Methods). Again paralleling the case of Neanderthal ancestry in modern humans^6^, introgressed anubis ancestry is most common in regions that are predicted to be least affected by linked selection (Fig. 2C; Spearman’s rho = 0.168, p = 1.73 × 10^−66^; Supplementary Methods). This pattern is driven by both genes and regulatory elements. Specifically, anubis ancestry per individual (n=442 Amboseli animals) is reduced, on average, by 7.03% in protein-coding regions relative to random, size-matched regions of the genome (±4.20% s.d.). The comparable reductions in promoters and putative peripheral blood mononuclear cell enhancers were 5.56% (±4.10% s.d.) and 6.22% (±4.20%), respectively.

Third, we tested whether introgressed anubis ancestry is positively correlated with local recombination rate^32^. This relationship is predicted under a model in which recombination influences the rate at which natural selection eliminates deleterious introgressed ancestry and uncouples deleterious from neutral introgressed variants (allowing neutral ancestry to persist). This prediction, documented across diverse taxa^4,38–41^, is also upheld in baboons (Fig. 2E; Spearman’s rho = 0.127, p = 1.48 × 10^−38^; Supplementary Methods). Strikingly, the magnitude of the observed correlation is similar to that reported for both Neanderthal and Denisovan gene flow into modern humans (Fig. 2F; Spearman’s rho = 0.17 for Neanderthal and 0.14 for Denisovan ancestry)^4^, despite substantially higher levels of anubis ancestry in the Amboseli baboons. Although our recombination map is derived from anubis baboons^32^ (Supplementary Methods), our results are robust to large window sizes where the recombination landscape is likely to be well-conserved between species^42^ (Table S3).

To investigate these patterns further, we took advantage of the dynamic history of admixture within the Amboseli population. At the beginning of long-term monitoring in 1971, observers considered all Amboseli animals to be yellow baboons^43^. Phenotypically anubis and admixed animals were observed to immigrate into the population starting in 1982, and microsatellite-based estimates suggest a subsequent increase in the proportion of anubis ancestry in the population over the following decades^25,44^. We observe the same pattern in the whole-genome resequencing data, which reveal an increase of 11.8% anubis ancestry from 1971 (29.6%) to 2020 (41.4%), as well as systematically higher anubis ancestry in immigrant males (Fig. 3A). However, both individuals born before the recent wave of anubis immigration and animals who have no known anubis or anubis-like ancestors are also clearly admixed (Fig. 3B). Additionally, most immigrants are not heavily anubis, and one immigrant was among the most yellow-like individuals in our sample (78.8% genome-wide yellow ancestry), indicating ongoing gene flow from outside the hybrid zone for both parental taxa.

**Figure 3:**
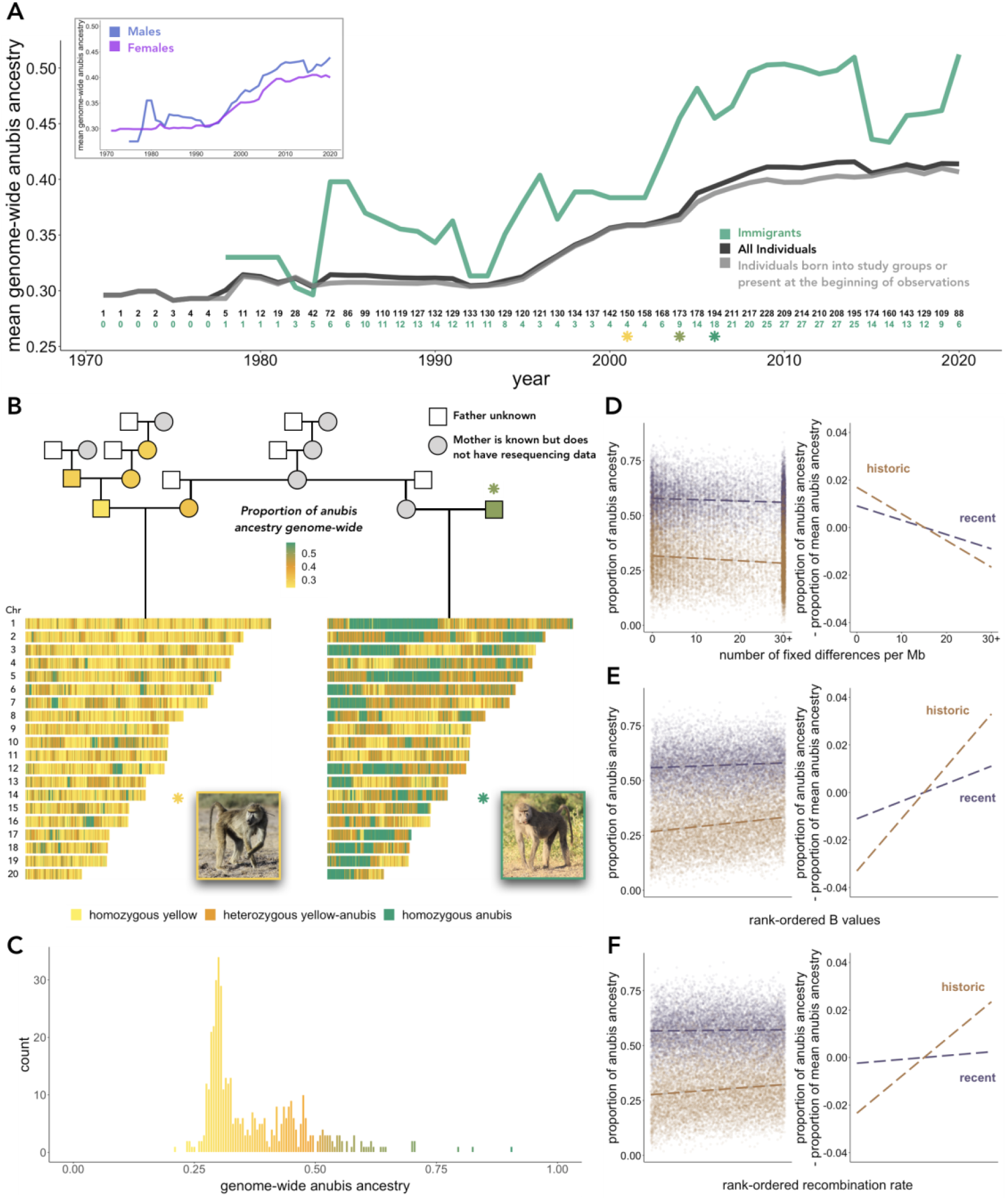
Recent and historic hybrid ancestry in Amboseli. **(A)** Mean genome-wide anubis ancestry in the Amboseli baboon population has increased since the late 1970s. Numbers above the x-axis indicate the number of individuals used to calculate mean anubis ancestry in the population each year (black numbers indicate all Amboseli individuals, green numbers indicate male immigrants). **(B)** Pedigree relationships and ancestry estimates for example Amboseli females of historic (left) and recent (right) hybrid ancestry. Pedigree individuals for whom we have resequencing data are filled by color according to their estimated global ancestry. The two focal females share the same maternal grandmother and were born only a few years apart (yellow and bright green asterisks in [A]). The father of the recent hybrid female was a phenotypic and genetic hybrid himself who immigrated in 2004 (olive green asterisk in [A]). **(C)** Histogram of genome-wide anubis ancestry in the population. The distribution is bimodal, roughly corresponding to historical versus recently admixed individuals. **(D-F)** Signatures of selection against introgression are more apparent for historical hybrids than recent hybrids. Panels display the relationship between introgressed anubis ancestry and (**D**) the rank-ordered number of fixed anubis baboon-yellow baboon differences, (**E**) mean B value, or (**F**) mean local recombination rate, across 250 kb windows of the genome. Right-hand panels facilitate comparison of slopes by mean-centering anubis ancestry within each data set to 0.

The Amboseli population today therefore consists of a mix of recently admixed animals and those unaffected by recent admixture, but who descend from admixture events that predate long-term monitoring^26^. By merging local ancestry calls with pedigree and near-daily observations of the same animals, we identify 188 “recently” admixed individuals whose pedigree includes a recent, anubis-like immigrant within the last 0-7 generations (mean=1.7 generations; note that due to previous gene flow, these animals’ genomes do not resemble classic F1, F2, or other early generation hybrids). We also classify 214 baboons as “historically” admixed, as they only contain anubis ancestry from before the start of long-term population monitoring (40 baboons could not be assigned to either hybrid class: Supplementary Methods). To estimate when historic admixture occurred, we assumed a single-pulse model of admixture in *DATES*^45^ and dated admixture in seven historical hybrids to between 18 and 665 generations ago (mean 283 ± 242 s.d.; Fig. S6). In contrast, gene flow in two more recently admixed individuals is dated to 5 and 21 generations ago. These ranges suggest repeated bouts of hybridization over hundreds of generations, including admixture events outside of the study population. Nevertheless, the composition of genome-wide anubis ancestry in the population is clearly bimodal (Fig. 3C), allowing us to evaluate the genomic distribution of introgressed anubis DNA before and after the first few hundred generations of selection. This is the period during which negative selection against introgression is thought to be strongest, including for Neanderthal introgression into humans^15–17^.

Stratifying individuals in the data set by admixture history reveals that signatures of selection against introgression are driven by historical admixture. Historically admixed individuals are more depleted of anubis ancestry in highly diverged and low B value regions of the genome than recently admixed individuals (Fig. 3D-E). Further, the relationship between introgressed anubis ancestry and recombination rate is exclusive to the historically admixed data set, even when investigating recombination rate at chromosome-level scales (Fig. 3F; Table S3; Supplementary Methods). The weaker signature of selection in recent hybrids likely reflects noise introduced by stochastic inheritance processes in the last few generations, long introgressed haplotypes, and recurring immigration by anubis-like animals into Amboseli. In contrast, sufficient generations have passed since historical admixture to break apart very large introgressed haplotypes, allowing us to observe non-random patterns of ancestry across the genome. This result emphasizes the importance of complementing field observations with genomic data, which provide insight into selective processes that operate over timescales longer than even the longest-running field studies.

## Selection against regulatory divergence

Analyses of human-Neanderthal admixture suggest that selection may have been particularly strong against regulatory variants^46^. If so, the introgressed regions that persist in modern humans have likely been purged of many alleles with large regulatory effects^7,47,48^. However, direct comparisons between the effect sizes of retained versus lost archaic alleles are difficult, as only a small fraction of archaic hominin ancestry still segregates in modern human genomes today^6,49^. Extant primate populations, where hybridization and selection are ongoing, provide a unique opportunity to test this hypothesis.

To test for selection against gene regulatory divergence in baboons, we paired genetic ancestry data with blood-derived RNA-sequencing data from 145 unique individuals (n=157 samples^50–52^; Table S1; Supplementary Methods). This data set includes whole blood collected from 2007 to 2010 and white blood cells collected from 2013 to 2018, which were processed and sequenced using distinct methods^50–52^. We therefore analyzed global and local ancestry effects on gene expression separately in these two data sets, controlling for age, sex, and kinship. Among 10,192 analyzed genes, we identified no significant associations between genome-wide ancestry and gene expression levels (10% FDR). In contrast, local genetic ancestry predicted gene expression levels for 20.1% (2,046) of tested genes in one or both data sets (Fig. 4A), and the estimated effects of local ancestry were highly concordant between data sets (Pearson’s R = 0.43, p < 10^−200^). In virtually all cases, local ancestry has an additive effect; we detected evidence for non-additive effects in only five genes, across both data sets (Supplementary Methods).

**Figure 4:**
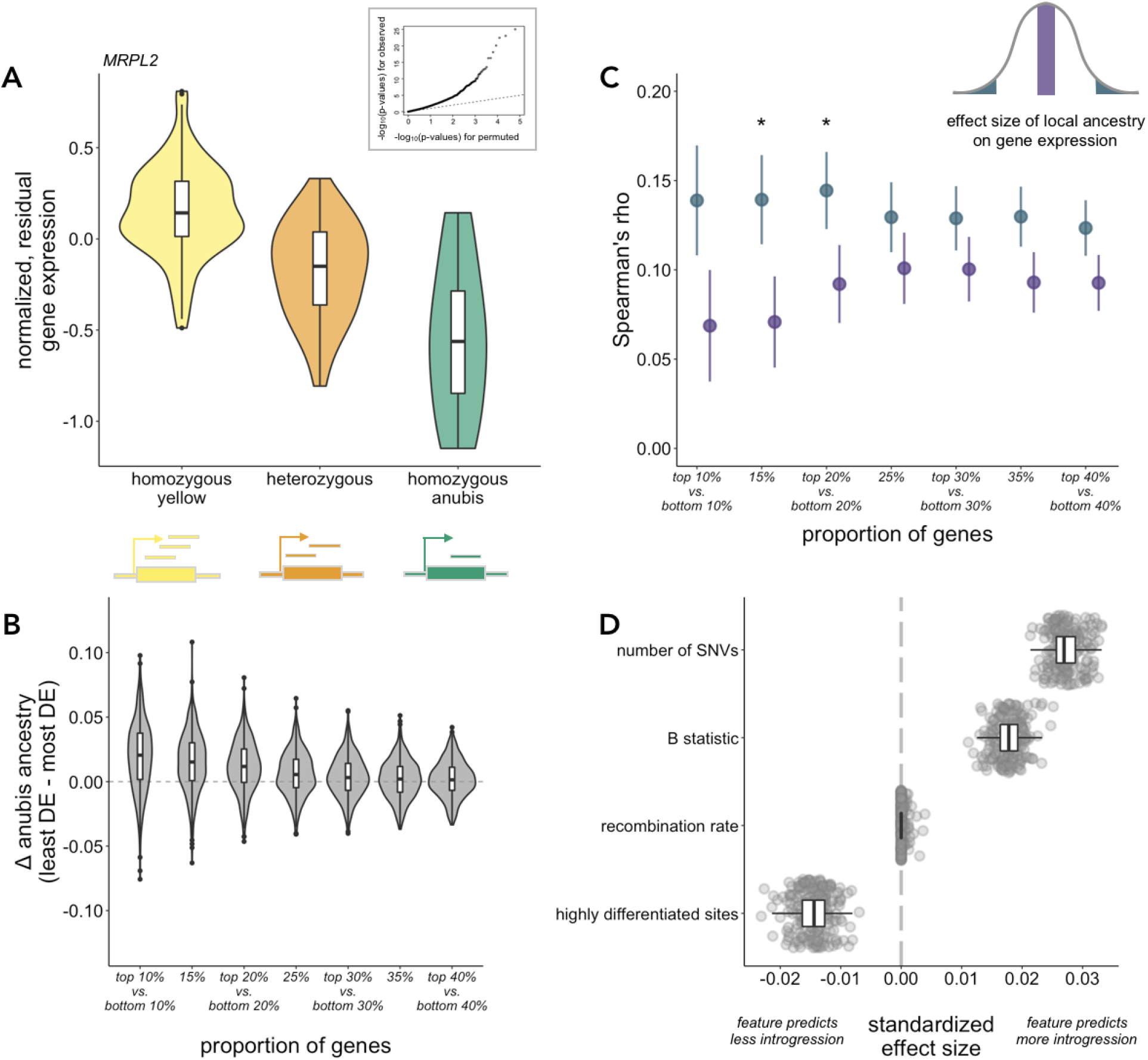
Selection against gene regulatory divergence and prediction of local introgression levels in the Amboseli baboons. **(A)** Local ancestry predicts gene expression in the Amboseli population, as depicted for an example gene (*MRPL2*) (*β* ∈ [−0.4,−0.34], local false sign rate < 10^−10^ in both gene expression datasets). Inset shows a quantile-quantile plot comparing p-value distributions from observed local ancestry effects on gene expression (y-axis) to permutation-based expectations under the null (x-axis). **(B)** Difference in introgressed anubis ancestry between genes with the smallest versus largest local ancestry effects on gene expression. Violin plots show the distribution of difference values per individual; boxplots show the median and inter-quartile range (all p < 0.05; Table S4). **(C)** Spearman’s rho between anubis introgression and recombination rate calculated for sets of genes with the largest (blue) versus smallest (purple) local ancestry absolute effect sizes. Asterisks denote bootstrapped p-value < 0.05 (Table S4) and error bars show standard deviations. **(D)** Distribution of effect sizes for features that consistently predict the extent of anubis introgression in Amboseli baboon genomes (Table S5). The number of single nucleotide variants (SNVs) is derived from unadmixed yellow and anubis baboon genomes (Supplementary Methods).

If introgressed alleles that perturb gene regulation are a primary target of selection, we reasoned that selection should purge anubis ancestry near genes where ancestry strongly affects gene expression. We therefore summarized evidence for local ancestry effects on gene expression at each gene as the mean effect size across the two gene expression data sets, after using multivariate adaptive shrinkage to refine effect size estimates^53^. The top 15% of genes with the strongest local ancestry effects on gene expression harbor, on average, 1.5% less anubis ancestry than those with the weakest local ancestry effects (Fig. 4B; paired t-test p = 1.10 × 10^−36^, n=442). This difference is exaggerated within historically admixed individuals (1.9% reduction; p = 1.26 × 10^−27^, n=214) (Table S4). We also compared the correlation between introgressed anubis ancestry and local recombination rate for matched quantiles of genes with the largest versus smallest local ancestry effects on gene expression. Spearman’s rho is significantly larger for genes with the largest effects of local genetic ancestry (Fig. 4C; Δ*_rho_* = 0.07 when comparing the most extreme 15% of genes; bootstrapped p-value < 0.05; Table S4). Combined with the depletion of introgressed sequence in regulatory elements, these results support the hypothesis that introgressed alleles affecting gene regulation are nonrandomly purged after hybridization. They therefore lend support to the idea that archaically introgressed variants with large effects on gene regulation are depleted in the genomes of modern humans today^7^.

## Predicting the genomic landscape of introgression

Finally, to investigate our ability to predict the genomic landscape of introgression, we modeled mean anubis ancestry as a function of local recombination rate, SNP density in unadmixed yellow and anubis populations, yellow-anubis genetic divergence, gene and enhancer content, linked selection, and local ancestry-associated gene expression in blood. To do so, we trained an elastic net regression model on non-overlapping 250 kb windows of the genome, representing 75% of the genome, and applied the model to a test set of windows in the remaining 25% of the genome (Supplementary Methods). We found that our predicted values were consistently positively correlated with observed levels of anubis ancestry in the test sets (mean Pearson’s R = 0.254 ± 0.0162 s.d., compared to 0.014 ± 0.011 s.d. for models fit to permuted data), with frequent contributions from features capturing local recombination rate, linked selection, genetic variation, and sequence divergence (Fig. 4D; Table S5). Further, when we stratified our data set based on the timing of admixture, we predicted introgressed anubis ancestry in historic hybrids significantly better than for recent hybrids (mean Pearson’s R = 0.265 ± 0.017 s.d. vs. 0.177 ± 0.018 s.d., bootstrapped p-value < 10^−3^). This difference likely arises because of the stronger relationship between historically introgressed ancestry and the sequence characteristics we highlight above (Fig. 3D-F).

Despite reduced predictive power in recent hybrids, our longitudinal data suggest that increases in anubis ancestry in Amboseli during the period of direct field observation are also non-randomly distributed throughout the genome. We used a linear fit to estimate the slope of change in anubis ancestry between 1979 and 2020 across 100 kb windows of the genome (n=25,797 windows). Controlling for the starting level of anubis ancestry in 1979 (p-value < 10^−300^), we found that windows with locally lower divergence (lower F_ST_) and higher recombination rates experienced larger increases in anubis ancestry during the past forty years, although both effect sizes are small (F_ST_ and recombination rate p-values <10^−3^, *β* = −2.965 × 10^−4^, 1.020 × 10^−4^, respectively; Table S6). B statistic values did not predict temporal change in anubis ancestry independently of recombination rate.

## Divergence and hybridization in primates

The behavioral and life history evidence to date in Amboseli—one of the largest and longest-running primate field sites in the world—indicates that hybrid baboons suffer no obvious fitness costs^27–29^. Nevertheless, genomic analysis reveals broad evidence for selection against admixture that is remarkably consistent with results obtained for archaic introgression in humans. They also support a hypothesis that can only be indirectly tested in hominins: that natural selection has acted to specifically eliminate introgressed alleles that substantially perturb gene regulation^7^. Our results identify subtle selection against hybridization that may help explain the maintenance of primate species boundaries in the face of frequent interspecific gene flow^1,54^.

The mode of selection against hybrids is still unclear. Unlike in humans, hybridization load is unlikely to explain our results: yellow and anubis baboons harbor similar levels of genetic diversity (<50% difference)^24,26^ compared to humans and Neanderthals, which exhibit more than three-fold difference^5,55^. Both hybrid incompatibilities and ecological selection, however, could play a role. Ecological selection is an attractive hypothesis because yellow baboons and anubis baboons are differentiated by pelage color and thickness; anubis baboons are thought to be associated with cooler, higher altitude environments in Kenya^56^; and high ambient temperatures in Amboseli predict elevated glucocorticoid levels in both sexes and reduced testosterone levels in males^57,58^. Indeed, baboon species appear to have moderately well-defined ecological niches on a pan-African scale, despite no clear correlation with ancestry in the region around Amboseli^59,60^. Assortative mating by ancestry—a known predictor of mating behavior in Amboseli—may also contribute to variation in the landscape of introgression^28^, although the responsible traits have not yet been identified. Understanding the genetic and phenotypic mechanisms that favor versus restrict interspecific gene flow in baboons, including the role of the X chromosome and adaptive introgression, therefore remains an important goal for future work.

Together, our findings illustrate the importance of contextualizing genomic data with phenotypic and demographic information to understand the evolutionary dynamics of admixture. Genomes harbor information about historical processes that stretch back many generations before the beginning of field observations, and can capture subtle signatures of selection that may not be obvious in natural populations where demographic stochasticity is high, sample sizes are modest, and the specific phenotypes under selection may be unknown. Conversely, field data reveal the range of phenotypic and fitness outcomes that are compatible with genomic signatures of selection. Indeed, genomic evidence alone has led some researchers to posit that the costs of modern human-archaic hominin interbreeding must have been high, perhaps reflecting species at the brink of reproductive incompatibility^61,62^. Our results point to the limits of these inferences by indicating that qualitatively similar evidence for selection against introgression can be compatible with primate hybrids that thrive^27–29,44^. Together, our results highlight the crucial role of other primates for understanding human evolution, especially for phenomena that are impossible to study in our lineage alone.

## Acknowledgements

We thank J. Altmann for her fundamental contributions to research on the Amboseli baboons, without which none of this work would have been possible. We also thank the University of Nairobi, Institute of Primate Research, the National Museums of Kenya, the members of the Amboseli-Longido pastoralist communities, the Enduimet Wildlife Management Area, Ker & Downey Safaris, Air Kenya, and Safarilink for their cooperation and assistance in the field. We thank members of the Amboseli Baboon Research Project long-term field team (S. Sayialel, G. Marinka, B. Oyath) and camp staff, and to T. Wango and V. Oudu for assistance in Nairobi. The baboon project database, Babase, was designed and programmed by K. Pinc and is expertly managed by N.H. Learn and J.B. Gordon. Our research was approved by the Kenya Wildlife Service (KWS), the Kenya National Environment Management Authority (NEMA), and the Kenya National Council for Science, Technology, and Innovation (NACOSTI). For a complete set of acknowledgments for the Amboseli Baboon Research Project, please visit http://amboselibaboons.nd.edu/acknowledgements/. We thank G. McVicker, P. Moorjani, K. Samuk, and members of the Tung lab, especially S. He, for helpful discussion and assistance with code. Finally, we thank L. Barreiro, A. Goldberg, P. Moorjani, and G. Coop and members of the Coop lab for constructive feedback on a previous version of this manuscript.

The Amboseli Baboon Research Project has been supported by the National Science Foundation and the National Institutes of Health, currently through NSF IOS 1456832, NIH R01AG053308, R01AG053330, R01HD088558, and P01AG031719, with additional support provided by Duke University, Princeton University, and the University of Notre Dame. This work was also supported by NSF BCS-1751783 to J.T. and T.P.V., BCS-2018897 to J.T. and A.S.F., and a Leakey Foundation Research Grant to T.P.V. We thank the North Carolina Biotechnology Center for support for high-performance computing resources (2016-IDG-1013). A.S.F. was supported by NSF GRFP (DGE #1644868) and NIH T32GM007754. Any opinions, findings, and conclusions expressed in this material are those of the authors and do not necessarily reflect the views of our funding bodies.

## Data Availability

All sequencing data generated for this study have been deposited in the NCBI Sequence Read Archive (SRA) under BioProject accession number PRJNA755322 (embargoed pending review). Previously generated genome resequencing data are available under PRJNA308870, PRJNA433868, PRJNA54005, PRJNA20425, PRJNA162517, PRJNA54003, PRJNA54009, PRJNA54007, PRJNA251424, and PRJNA295782, and previously generated RNA-seq data are deposited under PRJNA269070, PRJNA480672, and PRJNA731520. Local ancestry calls and masked genotype data have been deposited on Zenodo (doi://10.5281/zenodo.5199534) (embargoed pending review).

## Code Availability

Scripts for all data analyses and figure generation are available at https://github.com/TaurVil/VilgalysFogel_Amboseli_admixture/.

## Contributions

T.P.V., A.S.F., and J.T. conceived and designed the study. A.S.F., J.A.A., R.S.M., J.K.W., L.S., S. K., T.N.V., E.A.A., S.C.A, and J.T. collected data. J.D.W. contributed data analysis tools. T. P.V., A.S.F., J.A.A., and J.A.R. processed data. T.P.V. and A.S.F. performed analyses. T.P.V., A.S.F., and J.T. interpreted analyses. T.P.V., A.S.F., and J.T. wrote the manuscript with edits and revisions from all other co-authors.

