## Supplementary Information for "Selection against admixture and gene regulatory divergence in a long-term primate field study"

### TABLE OF CONTENTS

***1. Study populations*** **4**

*1.1. The Amboseli baboon population* 4

*1.2. Other anubis and yellow baboon populations* 4

**2. Sample collection 4**

**3. Data generation 5**

**4. Mapping and variant calling 5**

**5. Resequencing data set: quality control for individual identity 6**

**6. Local ancestry calling 7**

*6.1. Statistical method* 7

*6.2. Reference panels* 7

*6.3. Local ancestry tracts* 7

**7. Accuracy of local ancestry assignment and comparisons to alternative methods 8**

*7.1. Simulated data and LCLAE optimization* 8

*7.2. Comparison to other methods* 9

*7.3. Consistency of ancestry calls within pedigree trios* 10

**8. $F_4$ statistics 11**

**9. Assessment of admixture outside of Amboseli 12**

*9.1. Principal components analysis and IBDmix* 12

*9.2. Identifying introgressed ancestry using LCLAE confirms admixed status of SNPRC yellow founders* 14

**10. Timing of admixture and ancestry tract lengths 14**

**11. Evidence for selection against anubis introgression 15**

*11.1. Introgressed ancestry as a function of divergence between parental taxa* 15

*11.2. Introgressed ancestry as a function of estimated background selection* 16

*11.3. Introgressed ancestry as a function of local recombination rates* 16

*11.4. Depletion of anubis ancestry in putatively functional regions of the genome* 17

**Code is available at https://github.com/TaurVil/VilgalysFogel_Amboseli_admixture.**

**List of Supplementary Figures and Tables**

**Figures**

- Figure S1. Quality control for individual identity in the Amboseli resequencing data.
- Figure S2. Accuracy of local ancestry assignment.
- Figure S3. Local ancestry call consistency within pedigree trios.
- Figure S4. The structure of genetic variation between yellow, anubis, and hybrid baboons.
- Figure S5. Admixture in yellow and anubis baboons outside of Amboseli.
- Figure S6. Estimated timing of admixture in the Amboseli baboons.
- Figure S7. Introgressed anubis ancestry is reduced in highly diverged regions of the genome.

**Tables**

- Table S1: Sample metadata
- Table S2: Evidence of selection against Neanderthal introgression into modern humans
- Table S3: Evidence of selection against anubis introgression in the Amboseli baboon population
- Table S4: Signatures of selection near genes with larger effects of local ancestry on gene expression
- Table S5: *glmnet* features and mean effect sizes for predicting introgressed anubis ancestry levels across the baboon genome
- Table S6: Results from a linear regression model predicting rate of anubis ancestry change over time
- Table S7: Results from a linear regression model predicting the proportion of pedigree

trio local ancestry inconsistencies

- Table S8: Mean estimated identity by descent between yellow baboons and anubis, Guinea, and hamadryas baboons via IBDmix
- Table S9: Mean estimated identity by descent between anubis baboons and yellow, Kinda, and chacma baboons via IBDmix

### *1. Study populations*

#### *1.1. The Amboseli baboon population*

This study primarily focuses on a wild population of baboons inhabiting the Amboseli ecosystem of southern Kenya. Individuals from this population have been subject to near daily monitoring by the Amboseli Baboon Research Project (ABRP) since 1971[1]. When long-term observation began, the population was thought to include only yellow baboons[2], but immigrant anubis baboons and yellow-anubis hybrids have been documented since 1982[3-5].

Within the present sample of Amboseli individuals, 42.5% are known descendants of these anubis/anubis-like immigrants based on pedigree evidence (hereafter, referred to as "recently admixed" individuals, n=188). As suggested previously and confirmed here, however, even those individuals with no recent anubis ancestry appear to be admixed from events that predate long-term monitoring (hereafter, referred to as "historically admixed" individuals; n=214)[6]. Recently admixed individuals therefore contain anubis ancestry from both recent and historical admixture events. For the 40 remaining Amboseli animals, classification into these two categories was ambiguous because animals had no recent anubis ancestor but elevated levels of anubis ancestry consistent with an undetected, recent anubis ancestor (mean ancestry ≥ 30%, n=26) or because we were not completely confident about the individual's identity (n=14, see Section 5).

In cases in which we analyzed the genomes of recently admixed and historically admixed individuals separately, we focused on chromosomes with ≥ 30% estimated anubis ancestry for the recently admixed group. We did so because anubis ancestry is distributed unevenly across the genomes for recently admixed individuals, who are products of historical admixture as well. Consequently, some chromosomes in recently admixed individuals primarily reflect historical, not recent, admixture. Despite this step, we note that the signatures we observe in recently admixed individuals are likely driven in part by historical ancestry that co-exists on chromosomes with recent anubis ancestry.

#### *1.2. Other anubis and yellow baboon populations*

To characterize the genetic structure of the yellow-anubis hybrid zone, we also analyzed individuals from two putatively unadmixed yellow baboon populations and four putatively unadmixed anubis baboon populations. Yellow baboons were sampled from (i) Mikumi National Park in central Tanzania (n=15); and (ii) the wild-caught founders of the Southwest National Primate Research Center (SNPRC, now part of the Texas Biomedical Research Institute), originally captured in Kenya in 1972 (n=7). Anubis baboons were sampled from: (i) the Aberdares range in central Kenya (n=2); (ii) the Maasai Mara National Reserve in Kenya (n=7); (iii) the wild-caught founders of the SNPRC colony (n=24); and (iv) the Washington National Primate Research Center (WNPRC; n=6). Because the trapping locations for the SNPRC founders were close to the known hybrid zone[6] (Fig. 1A), it is possible that the colony founders may have been admixed themselves. This possibility was supported in our analyses for the yellow baboon founders, leading us to mask potentially introgressed regions when calculating reference population allele frequencies (Fig. 1; Section 6; Section 9). All other populations are far from the hybrid zone and unlikely to be admixed based on genetic data, phenotypic data, and/or field reports[6-10].

### 2. Sample collection

In this study, we used a combination of previously published and newly generated genomic data. Previously published whole-genome resequencing data were available for baboons from Amboseli (n=22), the Aberdares (n=2), Maasai Mara (n=7), Mikumi (n=5), the SNPRC (n=33), and the WNPRC (n=6)[6,8,11,12]. Previously published RNA-seq data were available for 145 unique individuals (157 samples) from Amboseli[13-15].

We also generated new whole-genome resequencing from individuals in Amboseli (n=420) using blood or other tissue samples collected by the Amboseli Baboon Research Project, for a total of 442 sequenced Amboseli baboons. The majority of samples were blood-derived, collected from animals that were anesthetized with a Telazol-loaded dart using a handheld blowgun. This procedure follows a highly conservative protocol focused on maintaining habituation and minimizing the possibility of injury[16,17]. Sample storage for DNA extraction varied over time because blood samples were collected from 1989-2018. However, in all cases, blood or purified white blood cell samples were placed in a storage medium (either a Tris-based storage buffer or RNALater) and archived at -80° C upon arrival in the United States. Samples for RNA extraction were collected in either PaxGene RNA tubes (Qiagen; for samples collected from 2007-2010[14]) or TruCulture "null" (media-only) tubes (Myriad RBM; for samples collected from 2013-2018[13,15]). Following sample collection, animals were placed in a covered holding cage to recover from the effects of the anesthesia and then released to their social group under observer monitoring. For an additional 32 samples used to generate resequencing data, we opportunistically collected non-blood tissue samples (e.g., from recovered corpses), stored in either 95% ethanol or RNALater until DNA extraction. An additional 5 samples were obtained from previously extracted DNA where the original sample was unavailable, but almost certain to be blood collected during darting.

### 3. Data generation

We extracted DNA for and sequenced 430 baboon samples. For 423 individuals (n=413 Amboseli animals, 10 Mikumi animals), we prepared DNA sequencing libraries using the NEBNext® Ultra™ II FS DNA Library Prep Kit for Illumina (New England Biolabs). We input 5 ng of DNA per library, and used half-reactions for fragmentation/end repair, adapter ligation, and PCR enrichment. Libraries were dual-indexed for sequencing using NEBNext® Multiplex Oligos for Illumina® (Dual Index Primers Sets 1 and 2) and amplified for 8 PCR cycles. We assessed library size and quality on a 2100 Bioanalyzer System (Agilent). For seven additional Amboseli individuals, DNA sequencing libraries were prepared by the Garvan Institute of Medical Research in Australia.

Libraries were sequenced on the Illumina HiSeq X Ten at MedGenome, Inc. or at the Garvan Institute. We generated moderate to high coverage resequencing data for seven Amboseli individuals (mean coverage from non-read duplicate reads mapped at MAPQ $\geq$ 10 = 35.56 $\pm$ 3.81 s.d.) and 10 yellow baboons from Mikumi (mean coverage = 13.38 $\pm$ 1.50 s.d.), and low coverage resequencing data for the remaining 413 individuals from Amboseli (mean coverage = 1.04 $\pm$ 0.19 s.d.). We combined these data with previously generated resequencing data from an additional 22 animals[6,12] to generate a dataset of 442 individuals from the Amboseli population.

### 4. Mapping and variant calling

We generated joint variant call format (VCF) files using a pipeline adapted from the Genome Analysis Toolkit (GATK[18-21]) Best Practices[22]. First, we used *TrimGalore*[23] to remove adapters and low-quality ends of reads (minimum Phred score threshold of 20) and imposed a minimum read length threshold of 50. Each sample was then processed individually to generate gVCF files as follows. We used *Picard*'s[24] FastqToSam to convert fastq files to BAM format and add unique read group names. We aligned reads to the *Panubis1.0* reference genome[25] with *bwa-mem*[26]. Specifically, we used piped commands for this step: Picard SamToFastq | bwa mem | Picard MergeBamAlignment. Next, we tagged PCR duplicates with *Picard*'s MarkDuplicates function. We then generated per-sample gVCF files with GATK *HaplotypeCaller*, using bases with minimum base quality scores of 10 and reads with minimum mapping quality scores of 20. Next, we performed joint genotyping using batches of gVCF files produced by GATK *CombineGVCFs*. Joint genotyping was performed with GATK *GenotypeGVCFs*. The resulting VCF files were then left-aligned using GATK *LeftAlignAndTrimVariants*, and alternate alleles that did not appear in any samples were removed with the *trimAlternates* option of the GATK *SelectVariants* tool.

We then filtered for high quality variants following GATK's recommended criteria for hard filtering for germline variants. Specifically, we retained biallelic variants with QualityByDepth (QD) > 2, Fisher strand bias (FS) < 60, root mean square mapping quality (MQ) > 40, mapping quality rank-sum test (MQRankSum) > -12.5, and rank sum test (ReadPosRankSum) > -8. To call a variant within the Amboseli baboon population, we also required that the position be genotyped in at least 20% of individuals and be genotyped in at least half of the anubis and yellow baboons used in our reference panels.

**5. Resequencing data set: quality control for individual identity**

For the Amboseli data set, we confirmed that our genetic data matched their donor labels based on correct assignment of sex and reconstruction of kin relationships consistent with the independently generated, multigenerational Amboseli pedigree. For cases in which RNA-seq data were also available, we also confirmed that genotypes from the resequencing data and the RNA-seq data matched within individuals.

For sex, we checked whether resequencing data from females had approximately 2x the proportion of mapped reads assigned to the X chromosome relative to resequencing data from males (Fig. S1A). For pedigree relatedness, we compared pedigree-based relatedness calculated using the R package *pedantics*[27] (version 1.7) to genetic estimates of relatedness based on the resequencing data, calculated using *lcMLkin*[28] (designed for low-coverage data). Pedigree maternal links are based on field observations of pregnancy and birth. Pedigree paternal links are based off demographic data (e.g., the presence of reproductively mature males in a female's social group at the time of conception) and microsatellite genotyping followed by paternity assignment by exclusion[29,30] (extragroup paternity has never been observed in this population).

For each individual, we plotted their pedigree versus resequencing-based estimates of relatedness with all other individuals in the data set, paying particular attention to concordant estimates for known close kin dyads (e.g., parent-offspring, full sibling, half-sibling, grandparent-grandoffspring) (Fig. S1B). Out of 442 individuals, we identified six cases where the sample did not match the original labeled individual and a further six individuals whose identity could not be unambiguously confirmed by the low-coverage data. In addition, two individuals were sampled from non-study groups (i.e., baboon groups that also live in the Amboseli ecosystem but are not regularly monitored to collect behavioral and demographic data). We removed these 14 individuals from analyses that depended on assignment of individual identity (i.e., because they required phenotypic or demographic data), but retained them for analyses that did not require individual-specific information.

For individuals with paired genome resequencing and RNA-seq data, we calculated the correlation between genotypes called from the resequencing data against genotypes called from all RNA-seq samples. In all cases, the highest resequencing-RNA-seq correlation was observed between samples from the same individual.

### 6. Local ancestry calling

#### *6.1. Statistical method*

We estimated local ancestry for each individual in the resequencing data set using LCLAE[6] (**L**ow **C**overage **L**ocal **A**ncestry **E**stimation). This method estimates the number of introgressed alleles at each SNP based on the genotype likelihoods for nearby sites (here, 0 = homozygous yellow; 1 = heterozygous; 2 = homozygous anubis). LCLAE thus draws on allele frequency differences and uncertain genotype calls integrated across variants to reliably assign ancestry from low coverage data. Specifically, LCLAE calculates the likelihood of each possible ancestry $Y_i$, where $i$ is 0, 1, or 2 and corresponds to the number of alleles of introgressed ancestry (here, anubis). This likelihood is given by:

$$lik(Y_i|G) = \sum_{j=0}^{2} \Pr\left(G_j\right) lik(Y_i|G)$$

where $G$ is the genotype (a set of three possible values) and $j$ refers to the number of alternate alleles. To combine information across SNPs, a composite likelihood is calculated by multiplying the probabilities of observed genotypes across sites in a predefined window size. The ancestry state with the greatest composite likelihood score is the one assigned for that SNP; in the case of ties, no ancestry is assigned. Then, for each SNP, a final ancestry call state is made using majority rule on all windows overlapping that SNP.

#### *6.2. Reference panels*

LCLAE requires allele frequency information for the parent taxa for inference. To generate this information, we used allele frequencies from the 7 SNPRC yellow baboon founders and the 24 SNPRC anubis baboon founders, all of whom were sequenced to high coverage[11]. We masked any regions of the genome that might be affected by introgression, as our analyses suggest that the SNPRC yellow baboon founders likely harbor low levels of anubis baboon ancestry from past admixture events (Fig. 1; Section 9).

To identify putative introgressed regions to mask for each reference panel individual, we used a conservative intersection set of loci identified as (i) introgressed based on local ancestry calls with LCLAE, using reference panel allele frequencies calculated without that individual included; and (ii) identity-by-descent (IBD) with members of the reference panel for the other species using IBDmix[31]. For IBDmix, which performs pairwise analyses, we required consensus IBD calls from at least 50% of reference animals from the other species. Nearly all potentially introgressed regions identified by LCLAE are included in the set of IBD calls identified by IBDmix (Section 9.2). Our approach masks 14.9-34.2% of SNPRC yellow founder genomes, and 2.0-6.6% of SNPRC anubis founder genomes. However, masking increases the density of ancestry-informative loci (i.e., SNPs with large allele frequency differences between yellow and anubis baboons) across the genome relative to unmasked genomes. It also improves ancestry calling consistency within Amboseli pedigree trios (Section 7.3).

#### *6.3. Local ancestry tracts*

We performed local ancestry analysis for all study subjects in all populations, including reference population animals after removing the focal animal from the reference panel before inference. To do so, we focused on ancestry informative markers that exhibited at least a 20% difference in allele frequency between the yellow and anubis reference populations and that were also genotyped in Amboseli. We calculated composite likelihood values for 35 kb windows centered on each ancestry informative marker and then estimated local ancestry at that marker using majority rule for sites within 35 kb upstream or downstream of the focal marker (i.e., ancestry was assigned based on the majority ancestry state when ≥50% of sites within the 70 kb centered on each site were assigned the same state). We removed ancestry calls from windows with fewer than 20 nearby ancestry informative markers, as low marker density contributes to errors in ancestry calling. These parameters and window sizes were selected based on simulations to maximize accurate ancestry assignment (Section 7.1). Finally, we collapsed sequential ancestry informative markers that were assigned the same ancestry state into contiguous tracts. We placed tract break points exactly halfway between sequential SNPs in which ancestry state assignments switched. We also removed ancestry tracts shorter than 1 kb, as simulations indicate that tracts of this size are unreliable (Section 7.1).

### 7. Accuracy of local ancestry assignment and comparisons to alternative methods

#### *7.1. Simulated data and LCLAE optimization*

To evaluate the accuracy of LCLAE local ancestry calls, we simulated 25 generations of admixture in a population of 1,000 baboons with 30% anubis ancestry (approximately the mean ancestry in the current population), discrete generations, limited immigration (10 individuals per generation), and a uniform recombination rate using SELAM[32]. Simulated individuals mated randomly and no variants were under selection. This simulation produced "true" local ancestry tracts for 25 individuals with a length distribution similar to those observed in real data from 9 Amboseli baboons sequenced at high coverage.

Variant-level genotypes for each simulated individual were drawn based on the simulated individual's known ancestry at each ancestry-informative marker and reference panel allele frequencies for yellow and anubis baboons. For these simulations, we used the same reference panels that we applied in real data analysis (7 yellow baboon and 24 anubis baboons from the SNPRC founders). We then simulated sequencing reads (10x coverage) for each individual using the program NEAT-genReads[33]. After mapping simulated reads using *bowtie2*[34] and calling sample genotypes as above, we called local ancestry for each simulated individual using LCLAE and the same SNPRC founder reference panels. To assess the effects of sequencing depth, we also down-sampled the mapped bam files to 1x coverage (similar to most of our resequencing data for Amboseli), re-called genotypes, and re-estimated local ancestry. To optimize the set of ancestry-informative sites and window sizes used to calculate composite likelihoods, we also explored the relative accuracy of local ancestry calls when setting the minimum yellow-anubis allele frequency difference for ancestry-informative sites to values between 5 and 50%, and using ancestry informative sites in window sizes of 10 to 50 kb, with windows twice that size used to smooth ancestry calls using majority rule.

To assess local ancestry calling accuracy, we calculated the percentage of the entire genome where the estimated ancestry matched the simulated ("true") ancestry. LCLAE performs very well for both high and low coverage data, correctly assigning between 93.1 and 97.3% of the genome to the appropriate ancestry state (Fig. S2A; mean accuracy = 94.8 ± 1.0% s.d. for high coverage sequencing and 94.6 ± 1.1% for 1x coverage sequencing). About half of the errors in calling local ancestry in the low coverage data were also errors in the high coverage data, suggesting that errors are often due to LCLAE's assumptions themselves, not to low coverage.

For window size, there is a trade-off between the number of ancestry informative sites used to make ancestry calls — there are more sites in larger windows — and the possibility of missing small ancestry tracts because larger windows can span multiple smaller tracts. For low (1x) coverage data, a 35 kb window size maximized the proportion of the genome that was assigned to the "true" ancestry state (96.9%; Fig. S2B). High coverage data follow a similar pattern, but tend to produce reasonably accurate estimates even with smaller window sizes, presumably due to a greater density of ancestry informative sites. Because most of our sequencing data for Amboseli is low coverage, we therefore used 35 kb windows in our subsequent analyses. We note that local ancestry calling is highly accurate for longer ancestry tracts (reaching 95% accuracy for tracts greater than 50 kb, even in low coverage data) but highly error-prone for tracts shorter than 1 kb (only 40% accuracy for high coverage data) (Fig. S2C-D). We therefore eliminated any tract calls less than 1 kb. We found no appreciable difference in the accuracy of ancestry assignments with minimum allele frequency differences between 5 and 30%. Accuracy decreases when using a higher threshold for yellow-anubis allele frequency differences (e.g., 50%) because of the sparsity of highly divergent sites with genotype calls, particularly for low coverage data. However, small differences in allele frequency are likely to be uninformative regarding ancestry in real data because estimated allele frequencies are drawn with sampling error and may not accurately reflect parental populations. We therefore chose a threshold of 20% minimum allele frequency difference between yellow and anubis baboons, which produces ancestry calls that are robust to sampling error in the reference populations (see below) and likely to represent true differences between species.

Finally, we assessed the robustness of our approach to errors in estimating reference panel allele frequencies (a concern because our panels were relatively small). To do so, we treated the estimated anubis and yellow baboon allele frequencies from the reference panel as "true" values, sampled genotypes based on these frequencies, and simulated reads for 10 yellow baboon and 10 anubis baboon genomes containing the sampled genotypes. Note that because we sampled genotypes from all common variants, the set of highly differentiated variants used for ancestry calling is different in the simulated data relative to the observed data. 78% of the ancestry informative sites identified in the original, observed data are also selected as ancestry informative sites in the simulated data (i.e., exhibit at least a 20% allele frequency difference between resampled, simulated anubis and yellow individuals). Of ancestry informative sites selected in the simulated data, 86.6% were also differentiated by at least a 20% yellow-anubis allele frequency difference in the original, observed data. Most importantly, substituting the allele frequencies estimated from these simulated reference panels for the original, observed allele frequencies had little effect on the accuracy of local ancestry calling. Ancestry calls using LCLAE for low coverage data still called the true, simulated ancestry state for 94.5% of the genome (compared to 94.6% when using the original, observed allele frequency estimates).

*7.2. Comparison to other methods*

In addition to estimating local ancestry using LCLAE, we tested two alternative approaches to calling local ancestry for low coverage data: Ancestry HMM[35] and AD-LIBS[36]. Whereas LCLAE uses a composite-likelihood method to call local ancestry, the other two approaches use Hidden Markov Models to infer the location of ancestry tract breaks. Further, while LCLAE works with genotype likelihoods, Ancestry HMM uses read counts from genotype calls and AD-LIBS uses mapped reads to create a pseudo-haploid genome per individual (a genome where each position is drawn at random from among mapped reads, if any are available). As for LCLAE, we focused on ancestry informative sites with a minimum allele frequency difference >20% between yellow and anubis baboons and placed tract breaks exactly in the middle between variants that were assigned different ancestry states. In these approaches, there is no analog to the window size parameter for LCLAE. For Ancestry HMM, we assumed 30% anubis ancestry and two pulses of anubis introgression, which were allowed to vary in time and degree of gene flow between species. For AD-LIBS, we also assumed 30% anubis ancestry.

For 10x coverage data, the Ancestry HMM approach performed best, matching the simulated ancestry for more than 99% of the genome (mean accuracy = 99.7 ± 0.1% s.d.; Fig. S2A). LCLAE performed slightly worse on 10x coverage (94.8 ± 1.0%). However, at low-coverage (1x), LCLAE retained high accuracy (94.6 ± 1.1%), whereas Ancestry HMM's accuracy dropped to 88.4 ± 3.8%. AD-LIBS performed surprisingly poorly at both 1x and 10x coverage, only assigning the correct ancestry 40 ± 5.6% of the time (Fig. S2A). This is likely because AD-LIBS uses only one mapped read per position, making it difficult to accurately assign ancestry between taxa that share substantial amounts of genetic variation (such as yellow and anubis baboons), as opposed to taxa with many fixed differences (e.g., polar and brown bears, the admixture case that motivated the original development of AD-LIBS[36]). Because most of our data set for this study is low-coverage data, we therefore chose to use LCLAE to assign local ancestry.

*7.3. Consistency of ancestry calls within pedigree trios*

We also evaluated the quality of our local ancestry calls using known parent-offspring trios from the Amboseli population. As a result of long-term monitoring, maternal-offspring links are known based on near-daily observations of pregnancies and the appearance of neonates; paternities have been assigned for the past several decades based on multilocus microsatellite genotyping[29,30]. Our resequencing data set contained 92 unique mother-father-offspring trios (n=181 unique low coverage individuals), in which, at any given locus, only a subset of possible local ancestry state calls would be consistent with the trio structure. We assessed consistency of local ancestry calls based on 73,975 loci across the genome that were spaced at least 35 kb apart and at least 50 kb from the ends of chromosomes. For each trio, we also removed loci that fell in tracts <1 kb length or that fell on the exact base pair that transitioned between ancestry states. For each trio at each remaining site, we evaluated whether the inferred ancestry of the offspring was consistent with the inferred ancestries of the parents (Fig. S3A-C). We then calculated the per-site proportion of ancestry state inconsistencies across all trios with data available for all three individuals.

Compared to the low-coverage reference panel used by Wall et al.[6], supplemented with 11 higher coverage Mikumi samples (5 of which were higher coverage data from individuals already sequenced by Wall et al.[6]), pedigree inconsistencies were lower when using high-coverage reference panels from the SNPRC anubis and yellow baboon founders (Fig. S3D; median proportion of pedigree inconsistencies using the Wall et al. panel = 0.283 ± 0.099 s.d. vs. 0.076 ± 0.062 s.d. using the SNPRC panel). Across loci, we achieved greater consistency when windows (i) contained more ancestry informative markers ($p<10^{-300}$); (ii) were more diverged between yellow and anubis baboons ($p<10^{-300}$; based on weighted $F_{ST}$ calculated using *vcftools*[37] [version 0.1.15]); and (iii) were in regions of elevated recombination ($p<10^{-32}$; mean recombination rates were calculated for 250 kb windows centered on each site and $\log_{10}$ transformed (see Section 11.3 for details on recombination rate estimates; Table S7). Across trios, inconsistencies were most common when one member of the trio was sequenced at particularly low coverage (Pearson's r = -0.553, p = 1.07 x $10^{-8}$; Fig. S3E).

**8. $F_4$ statistics**

In addition to estimating admixture using a local ancestry approach, we used $F_4$-ratio estimation[38-41] as an orthogonal method to estimate the anubis ancestry proportions of Amboseli individuals. Following the nomenclature of Patterson et al.[38], $F_4$-ratios estimate the ancestry proportion of members of population $X$ as a combination of two sources, populations $B'$ and $C'$. Because $B'$ and $C'$ are unknown ancestral populations, they are represented by modern populations $B$ and $C$. This method also requires two additional populations: population $A$, which has not contributed to population $X$ but is a sister group to population $B$, and an outgroup, $O$, to populations $A$, $B$, and $C$. The contribution of population $B'$ to population $X$ is α (where 1-α therefore corresponds to the contribution of population $C'$ to population $X$) and $\hat{\alpha} = \frac{f_4(A,O;X,C)}{f_4(A,O;B,C)}$ (see Patterson et al.[38] for details).

To estimate $F_4$ ratios, we focused on Amboseli animals for which we had high coverage data (n=9), which include two recently admixed animals and seven historically admixed animals. Amboseli is therefore population $X$. We also used data from the SNPRC anubis founders[11] (population $B$; n=24), Mikumi National Park in Tanzania (population $C$; n=10), hamadryas baboons and Guinea baboons resequenced by the Baboon Genome Sequencing Consortium[8] (as two alternatives for population $A$; n=2 in each case), and either the gelada monkey (also from the Baboon Genome Sequencing Consortium) or rhesus macaque as the outgroup $O$ (n=1 in each case). We mapped these data to the rhesus macaque genome (*MacaM*)[42] using *bowtie2*[34]. Duplicate reads were removed using the *MarkDuplicates* function from *Picard*[24] and reads with MAPQ less than 10 were removed using *samtools*[43] *view*. gVCF files were generated using GATK *HaplotypeCaller*[21] requiring a minimum base quality ≥ 20 and then merged across individuals using *CombineGVCFs*. Genotypes were then called using *GenotypeGVCFs*. We filtered for high quality variants following GATK's recommended criteria for hard filtering for germline variants (see Section 4) and retained biallelic SNPs that were typed within all individuals in the sample. We removed indels, singleton and doubleton variants, and removed clusters of 3 or more variants that fell within a 10 bp window, as dense clusters of SNPs are enriched for false calls. Since we used the macaque as one possible outgroup, we added the macaque genotype (homozygous reference) as an additional sample at all sites.

We next converted our calls, in vcf format, to PLINK format (using *vcftools*[37], *bcftools*[44], and *PLINK*[45-47] [version v1.90b3.36]) and then to EIGENSTRAT format (using *convertf*[38] in the *EIGENSOFT* package [version 6.1.4]). To calculate the $F_4$ ratio, we input the EIGENSTRAT formatted data into *admixr*[48] (version 0.9.1), an R package that builds on the *ADMIXTOOLS*[38] software suite. Using the *f4ratio* function*,* we generated estimates of α per chromosome separately for recently and historically admixed Amboseli animals (i.e., we treated them as two separate $X$ populations) for the following phylogenetic configurations ($A$, $B$, $C$, $O$): (i) hamadryas, SNPRC anubis, Mikumi yellow, macaque; (ii) Guinea, SNPRC anubis, Mikumi yellow, macaque; (iii) hamadryas, SNPRC anubis, Mikumi yellow, gelada; and (iv) Guinea, SNPRC anubis, Mikumi yellow, gelada. To obtain a genome-wide estimate of α, we first averaged α estimates for the different phylogenetic configurations per chromosome. Estimates of α were highly stable across phylogenetic configurations (range of s.d. across phylogenetic configurations per chromosome = 0.004-0.037). We then averaged values of α across chromosomes, weighted by chromosome length. Using this method, the genome-wide proportion of recently admixed Amboseli genomes derived from anubis gene flow is 0.508 while for historically admixed animals it is 0.232.

### 9. Assessment of admixture outside of Amboseli

#### *9.1. Principal components analysis and IBDmix*

Yellow-anubis admixture is well-documented within the Amboseli population, but may also occur outside of the known hybrid zone (as suggested by identification of an admixed animal from the Aberdares region of Kenya[8]). Understanding the extent of admixture in this region is important for placing admixture in Amboseli in context. In addition, understanding the extent of admixture outside the known hybrid zone is crucial for accurately defining reference panels when calling local ancestry.

To investigate the structure of our sample, we first used principal components analysis (PCA) to visualize the major axes of variation among high-coverage genomes from Amboseli (n=9), the Aberdares region (n=2), Mikumi National Park in Tanzania (n=11), the SNPRC founders (n=31), and 2 additional SNPRC anubis baboons from the Baboon Genome Project (Fig. 1B)[8,11]. We also performed a second PCA on the same set of individuals, plus animals sequenced at low coverage from Mikumi (n=4), the WNPRC (n=6), and Maasai Mara National Reserve in Kenya (n=7)[6] (Fig. S4). As expected, in both cases PC1 captures the majority of the variance in the genotype data (84% and 53% proportion variance explained in high and all-coverage genomes, respectively) and separates yellow and anubis baboons, with recently admixed individuals lying between the two extremes. When including a mix of high and low coverage data, samples also separate based on sequencing depth on PC2 and, to a lesser extent, on PC1 (Fig. S4). Interestingly, however, yellow baboon founders from the SNPRC cluster closely with Amboseli baboons affected by historical admixture (Fig. 1B).

The results of the PCA suggest that founders of the SNPRC colony that were labeled as yellow may contain nonnegligible amounts of ancestry from anubis baboons; indeed, these animals were trapped relatively close to the known hybrid zone, near Kibwezi and Darajani (Fig. 1A). We therefore tested whether we could detect evidence of anubis ancestry by identifying regions of the genome that are identical by descent (IBD) between the SNPRC yellow baboon founders and anubis baboons. To do so, we used IBDmix, a method designed to identify segments in a sample of test individuals (e.g., modern humans, in the original application) that are IBD with a candidate source genome (e.g., Neanderthal in the original application)[31]. Importantly, this method avoids assuming that members of a given population are unadmixed *a priori* (as has commonly been assumed, for example, for Neanderthal ancestry in human populations of African ancestry). In addition to assessing IBD with anubis baboons among SNPRC yellow baboon founders (n=7), we ran IBDmix on SNPRC anubis baboon founders (n=24) and 2 additional anubis colony members, yellow baboons from Mikumi (n=11), and anubis baboons from the Aberdares (n=2).

To assess IBDmix's false positive rates due to either misassignment or incomplete lineage sorting, we also evaluated IBD of yellow baboons with hamadryas baboons (n=2) and Guinea baboons (n=2), which share a divergence date with anubis baboons (i.e., all are "northern clade" lineages that separated from the ancestors of yellow baboons ~1.4 mya[8]). There are no known records of wild yellow-Guinea or yellow-hamadryas hybridization in the literature, and the current geographic ranges of these species makes gene flow implausible (Fig. 1A). Similarly, we used data from Kinda baboons (n=3) and chacma baboons (n=2)—two "southern clade" baboon lineages that share a divergence date with yellow baboons—to assess misassignment of yellow baboon IBD segments in anubis baboon populations.

We focused on high quality genotype data for samples analyzed with IBDmix. Specifically, we used the same filtering strategy as in Section 8 ($F_4$ statistics), except applied to the baboon genome, which retained 28.2 million variants. We then ran IBDmix with default settings, except that we decreased the source individual genotyping error rate to match IBDmix's settings for modern samples (the default settings assume that the source is an archaic genome, whereas all of our samples come from high-coverage, modern samples). We repeated this procedure for each test sample, across all possible source samples, to produce a set of regions that were identified as IBD for each test sample compared to each source sample. We focused on IBD regions that were at least 50 kb long and supported by a LOD score greater than 10 (stringent criteria that are likely to strongly enrich for true shared ancestry[31]). We then calculated the mean proportion of each test sample that was inferred to be IBD across possible source individuals. Notably, IBDmix does not differentiate between heterozygous states in which one allele is IBD with the source sample versus homozygous states in which both alleles are IBD. Thus, the proportion IBD reflects the proportion of the genome in which one *or* both alleles (but most likely one) is IBD between test and source individuals.

In yellow baboons from Mikumi, which is far from the known hybrid zone (Fig. 1A), a mean of 5.7 ± 0.8% (s.d.) was estimated to be IBD with anubis baboons, depending on the test and source individuals. This value is likely an overestimate, reflecting incomplete lineage sorting and erroneous calls as well as true shared ancestry. Indeed, 0.9 ± 0.1% of genomes from the same individuals were estimated to be IBD with Guinea baboons and 3.2 ± 0.3% were estimated to be IBD with hamadryas baboons. Given that the range of Guinea baboons is over 5,000 km from Mikumi, we view the most common explanation for IBD calls to be error or ILS. A similar pattern was observed in anubis baboons from the SNPRC and one of the two anubis baboons from the Aberdares: low levels of IBD with yellow baboons were estimated genome-wide (5.0 ± 0.8%), but the ranges of these values did not greatly exceed estimates for IBD with other southern clade baboons (geographically distant chacma and Kinda baboons) (Table S8). The second Aberdares anubis baboon was previously identified as an admixed individual by the Baboon Genome Consortium, which reported ~546 Mb of yellow baboon-derived nuclear DNA[8]. Our analysis supports that finding, with an IBD estimate for one or both alleles for 15.2% of the genome.

In contrast to the other non-Amboseli populations, yellow baboon founders from the SNPRC colony clearly appear to be admixed. IBDmix estimates an average IBD of 26.5 ± 6.7% for SNPRC yellow founders with anubis source individuals, in contrast to 5.7 ± 0.8% for Mikumi yellow baboons (t-test $p = 1.68 \times 10^{-4}$). Estimated IBD for SNPRC yellow founders with anubis baboons also exceeds the estimates for SNPRC yellow IBD with Guinea baboons or hamadryas baboons (vs. Guinea baboons: mean difference = 20%, paired t-test $p = 3.37 \times 10^{-5}$; vs. hamadryas baboons: 12%, $p = 7.16 \times 10^{-5}$). Further, when SNPRC yellow founders were inferred to be IBD against one anubis source individual, this inference tended to be highly consistent across other anubis source individuals, leading to extended "peaks" where IBD is broadly supported across most or all source animals. Because IBDmix is agnostic to the direction of gene flow, we therefore also observe a large fraction of anubis baboon genomes that appear to be IBD with yellow baboons when SNPRC yellow founders are used as source individuals (17.6-30.8%; Table S9). This value drops to 3.1-6.9% if Mikumi yellow baboons are used as source individuals.

*9.2. Identifying introgressed ancestry using LCLAE confirms admixed status of SNPRC yellow founders*

We complemented our IBDmix analyses by testing all yellow and anubis baboon samples for admixture using LCLAE, which (unlike IBDmix) distinguishes between heterozygous and homozygous ancestry states and allowed us to include low coverage samples. For each individual, we called local ancestry states using reference panels for anubis and yellow baboon allele frequencies composed of all yellow and anubis individuals except the focal individual. LCLAE assigned all high coverage samples from the SNPRC anubis founders and Mikumi yellows, and one of the anubis baboons sampled from the Aberdares, to their *a priori*-defined ancestries for >95% of the genome. Low coverage samples were similarly categorized, but low coverage appears to pull admixture proportions away from the extremes (i.e., in Mikumi, the mean proportion of anubis ancestry is 3.3 ± 0.2% (s.d.) in high coverage samples, but 14.3 ± 6.4% in low coverage samples). As expected based on IBDmix and PCA results, high coverage SNPRC yellow founders were estimated to have 17.1 ± 4.2% anubis ancestry, and the high coverage admixed anubis animal from the Aberdares was estimated to have 11.0% yellow baboon ancestry (Fig. S5A).

For comparison between LCLAE and IBDmix, we collapsed homozygous and heterozygous introgressed ancestry estimates from LCLAE because IBDmix does not differentiate between these two alternatives. Genome-wide estimates of introgressed (i.e., IBD) ancestry are highly correlated between LCLAE estimates and IBDmix estimates (Pearson's r > 0.99, p-value < $10^{-21}$ for Mikumi and SNPRC yellow baboons; r > 0.99, p-value < $10^{-20}$ for Aberdares and SNPRC anubis founders; Fig. S5B-C). IBDmix tends to detect more shared ancestry than LCLAE, although this effect lessens if IBD is only inferred for loci where the results for most (at least 70%) source individuals agree. On a site-specific level, regions identified using LCLAE were more than 20x as likely to be IBD based on IBDmix analysis across species, relative to the genomic background (i.e., randomly sampled, size-matched regions of the genome).

### 10. Timing of admixture and ancestry tract lengths

To estimate the timing of admixture in high coverage samples from Amboseli, we used DATES[49]. This method estimates the timing of admixture by fitting an exponential regression to the decay in ancestry covariance by genetic distance. We evaluated admixture timing in seven "historic" Amboseli hybrids and two "recent" Amboseli hybrids sequenced at medium to high coverage. In all cases, we considered a simple model of a single pulse of admixture. Although we believe that multiple waves of admixture have likely influenced baboon evolution in East Africa, current methods are not able to differentiate separate immigration events from the same source population, especially when the level of discontinuity in these immigration waves is unclear. Thus, we view the DATES estimates as composite estimates of the timing of anubis-yellow admixture across the genome, weighted towards more recent events.

To run DATES, we used the SNPRC anubis founders and Mikumi yellow baboons to represent the source of anubis and yellow ancestry for Amboseli baboons. After filtering for common variants called in high coverage Amboseli samples (minor allele frequency ≥ 5%), we converted SNP locations into a genetic map using baboon recombination rates[11] (Section 11.3). Specifically, we calculated the mean recombination rate per 100 kb, removed unreasonably large local recombination rates (>100x larger than the median recombination rate), and then scaled LDhelmet's population-scaled recombination rate to centiMorgans based on the baboon chromosome lengths estimated by Cox et al.[50]. We then fit a loess smoothing function to predict the position (in Morgans) of each SNP along the genome and used DATES to estimate the timing of admixture, using a maximum distance of 0.5 Morgans and a bin size of 0.001 Morgans.

Consistent with long-term field observations in Amboseli, the number of generations since admixture for recent hybrids (5 and 21 generations) is substantially smaller than for historic hybrids (Fig. S6A). This inference is also consistent with longer heterozygous and homozygous anubis ancestry tracts in individuals with known recent anubis ancestors, compared to Amboseli animals with no known recent anubis ancestors (Fig. 3B; Fig. S6B).

**11. Evidence for selection against anubis introgression**

To investigate whether signatures of selection against archaic introgression in humans are paralleled in baboons, we performed three tests. First, we asked whether regions of the genome that exhibit greater divergence between the parent species have reduced introgressed (anubis) ancestry in Amboseli. In humans, loci that contain fixed or near-fixed differences between humans and Neanderthals are more likely to be depleted of archaic ancestry[51]. Second, we tested whether regions of the genome that are predicted to be more affected by background selection also exhibit reduced introgressed ancestry, as reported for Neanderthal introgression into the human genome by Sankararaman et al.[52]. Third, we tested whether introgressed ancestry is depleted in regions of the genome with low mean recombination rate, as expected if deleterious alleles are selectively eliminated along with linked sequence. This pattern is also observed for Neanderthal and Denisovan ancestry in modern humans as well as in other hybridizing animal and plant taxa[53-56]. For all three analyses, we calculated mean anubis ancestry for non-overlapping windows of the genome (100-1000 kb) in (i) all baboons from Amboseli (n=442); (ii) historical hybrids only (n=214); and (iii) recent hybrids only (n=188) (see Section 1.1 for how recent and historical hybrids were defined). We report results from local ancestry calling with LCLAE in the main text. However, the results are qualitatively unchanged if we use ancestry assignments based on Ancestry HMM[35] (Table S3).

*11.1. Introgressed ancestry as a function of divergence between parental taxa*

Using all non-Amboseli yellow and anubis baboons (after masking loci potentially affected by introgression: Section 6.2), we counted the number of fixed differences that occurred in non-overlapping 250 kb windows across the genome. As reported in the main text ("Selection against introgression in Amboseli"; Fig. 2), we observed a strong negative relationship between introgressed ancestry in these windows and the number of fixed differences between yellow and anubis baboons (Spearman's rho = -0.119, p = $6.44 \times 10^{-31}$), which is largely driven by historical hybrids (Fig. 2A, 3D). As an alternative approach, we also identified highly differentiated sites based on allele frequencies estimated from anubis baboon and yellow baboon reference panel individuals (Section 6.2). We calculated $F_{ST}$ for every biallelic variant called in at least 5 anubis and 5 yellow baboons (n=21 million biallelic SNPs) as the loss in heterozygosity between species relative to a panmictic population[57]. Specifically, $F_{ST} = \frac{Ehet_{panmictic} - mean(Ehet_{populations})}{Ehet_{panmictic}}$, where *Ehet* (the expected heterozygosity) is calculated for each population and for a hypothetical panmictic population from observed allele frequencies, assuming Hardy-Weinberg equilibria. Introgressed ancestry is also negatively correlated with the number of highly differentiated sites (Fig. S7). Unsurprisingly, this relationship is strongest for nearly fixed differences ($F_{ST} \geq 0.75$, Spearman's rho = -0.109, p = $1.21 \times 10^{-28}$) and weaker, although still significant, for sites that are less differentiated (e.g., $F_{ST} \geq 0.5$; Spearman's rho = -

0.042, $p = 2.12 \times 10^{-5}$). For both fixed differences and highly differentiated sites, this pattern was robust across window sizes ranging from 100-1000 kb.

*11.2. Introgressed ancestry as a function of estimated background selection*

To estimate the degree of background selection across the *Panubis1.0* baboon genome[25], we used the B statistic from McVicker et al.[58]. B values are estimates of the proportion of nucleotide diversity maintained in the face of purifying selection, as a function of the deleterious mutation rate, the strength of selection, and recombination rate. These values assume that selection acts multiplicatively over loci and that the strength of selection is sufficient to ignore deleterious alleles in homozygous states[58]. B values near 0 represent strong background selection (i.e., near complete removal of diversity due to natural selection at linked sites), whereas large values correspond to little background selection. Following McVicker, et al. [58] and Sankararaman, et al. [52], we distinguished two classes of sites under selection: base pairs in exons (including UTRs) and base pairs in introns or promoter regions (defined as 10 kb upstream of gene transcription start sites). Exonic regions were assigned a deleterious mutation rate of $7.4 \times 10^{-8}$ mutations per base pair, and introns and promoters were assigned a deleterious mutation rate of $8.4 \times 10^{-10}$ mutations per base pair. Selection coefficients were drawn from an exponential distribution with mean $2.5 \times 10^{-3}$ and mean $1.0 \times 10^{-5}$ for exons and introns/promoters, respectively. We note that while the choice of these parameters is somewhat subjective, we used the same values as inferred for the human genome to maximize comparability; further, our analyses are focused on relative rather than absolute values of B. We note that, although our estimates are based on the anubis baboon reference genome, chromosomal structure and gene locations are highly conserved between baboon species.

As reported in the main text ("Selection against introgression in Amboseli"; Fig. 2), we observe reduced introgressed ancestry in 250 kb windows of the genome with higher levels of linked selection (lower B values). This result is stronger in historical hybrids than for those with recent anubis ancestry, and robust to changes in window size ranging from 100-1000 kb (Table S3). Similarly, we find that regions of the genome that contain more variable sites (suggesting weaker background selection) also have higher levels of introgressed ancestry (Spearman's rho = 0.196, $p = 4.83 \times 10^{-90}$ for 250 kb windows).

*11.3. Introgressed ancestry as a function of local recombination rates*

To evaluate the relationship between introgressed ancestry and local recombination rates, we required a fine-scale recombination map for baboons. An LD-based recombination map is available for the anubis baboon from Robinson et al.[11], but the published version is linked to an older version of the baboon genome (*Panu2.0*[8]) than the version we used here (*Panubis1.0*[25]). We therefore regenerated the anubis baboon recombination map using *LDhelmet*[59] (v1.9) and resequencing data from the 24 SNPRC anubis baboon founders[11]. We first phased the genomes using *Beagle*[60] (v5.0), after excluding uninformative singletons and pruning variants to at least 10 bp distance apart. We then ran *LDhelmet* with a block size penalty of 5 (as recommended by Chan et al.[59]), an $N_e = 40{,}000$ (based on[61]), and default parameter values except where noted. We set the population-scaled mutation rate, $\theta$, to 0.0016 in baboons, which is an approximation of the value of $\theta$ calculated from the number of variants in the input data prior to pruning. We used a window size of 50 and ran the Markov chain Monte Carlo inference for $1 \times 10^6$ iterations, following a burn-in period of $10^5$ iterations. The resulting estimates are in terms of $\rho$ ($= 4N_er$), the population-scaled recombination rate. We converted these values to centiMorgans based on the lengths of baboon chromosomes reported by Cox et al.[50].

As reported in the main text ("Selection against introgression in Amboseli"; Fig. 2), we observed increased introgressed ancestry in 250 kb windows of the genome with increased mean recombination rates. This result is also driven by historical hybrids rather than individuals with recent anubis ancestry, and robust to changes in window size ranging from 100-1000 kb (Table S3).

*11.4. Depletion of anubis ancestry in putatively functional regions of the genome*

We tested whether anubis ancestry was depleted in regions of the genome with known functional importance. We focused on coding sequences, gene promoters, and enhancers, in each case comparing anubis ancestry levels to size-matched, putatively neutral regions more than 100 kb away from annotated elements (i.e., genes, promoters, and enhancers). We used gene annotations from NCBI to define coding sequences[62], and defined promoters as the 10 kb upstream of the most 5' transcription start site. Because baboon enhancer annotations are not available, we defined putative baboon enhancers by projecting coordinates from ENCODE H3K4me1 ChIP-seq data for human peripheral blood mononuclear cells[63] onto the *Panubis1.0* genome using the UCSC Genome Browser liftover tool[64] and a liftover file available at https://github.com/TaurVil/VilgalysFogel_Amboseli_admixture/03_Resources/Resources/. We then identified five sets of size-matched regions using the shuffle command in *bedtools*[65] (version 2.25.0), excluding regions that were within 100 kb of annotated genes, promoters, or enhancers. For each type of element and each individual, we calculated the proportion of anubis ancestry and estimated mean anubis ancestry across all elements, yielding one estimate per individual. For each element type, we then compared these values to size-matched, putatively neutral regions using paired t-tests (n=442 individuals).

### 12. Comparative evidence from archaic hominin admixture

To provide context for our findings, we compared them to evidence for selection against archaic introgression in humans, focusing specifically on admixture with Neanderthals. All three of the tests we used have been previously reported in the literature[51-53], but using different human reference genomes and differing amounts of information (based on what was available at the time) on Neanderthal-human divergence and introgression. We therefore re-performed them here, which allowed us to follow an analysis pipeline that was as parallel to our analysis of the baboons as possible.

To do so, we first downloaded Neanderthal ancestry calls for HapMap CEU individuals from[66]. Following[53,66], we converted posterior probabilities for Neanderthal ancestry to 0/1 ancestry calls based on a threshold of 0.42 for Neanderthal ancestry (this approach assumes that homozygous Neanderthal tracts, i.e., ancestry state 2, are rare). We then averaged ancestry states over all sites in a given genomic window for each individual and averaged these per-individual values to obtain an estimate of introgressed (Neanderthal) ancestry in the 1000 Genomes CEU population[67]. Importantly, population-level introgressed archaic ancestry across the genome is highly correlated with the haplotype-based calls of Sankararaman et al.[52] (at the 50 kb scale, Spearman's $\rho = 0.88$; at the 500 kb scale, $\rho = 0.94$).

B values for the human genome were obtained from[58] and converted to hg19/gr37 coordinates using the Ensembl AssemblyConverter[68]. To calculate window-based recombination rates, we used the HapMap Project recombination maps for CEU individuals[69]. Finally, to identify fixed differences between modern humans and Neanderthals, we downloaded and merged genotype calls for 3 high coverage Neanderthals (the Altai Neanderthal[70], the Vindija Neanderthal[71], and the Chagyrskaya Neanderthal[72]) from[73]. We then identified fixed differences between Neanderthals and modern humans by filtering for single nucleotide variants that were called in at least 2 of the high coverage Neanderthal samples and fixed for the alternate allele in Neanderthals. We then removed variants that were variable in African populations in the 1000 Genomes Project[67]. This left 769,488 variants that appear to be fixed differences between Neanderthal and modern human genomes, and are at the very least highly differentiated between modern humans and Neanderthals.

We calculated the proportion of Neanderthal ancestry for the CEU population for non-overlapping 250 kb windows across the genome. We excluded windows in which no alleles were called for at least two of the three Neanderthal genomes. These proportions are negatively correlated with the number of fixed differences, positively correlated with B values (i.e., reduced background selection), and positively correlated with local recombination rate, as expected based on previous reports[51-53] (Fig. 2; Table S2).

### 13. Ancestry-associated gene expression levels

*13.1. Data processing*

To investigate the contribution of genetic ancestry to gene expression, as well as the relationship between ancestry effects and signatures of selection, we used RNA-seq gene expression data from 157 samples, representing 145 unique individuals. Data from 63 samples were obtained from whole blood and reported in[14]; data from 94 samples were obtained from white blood cells and reported in[13,15] (Table S1). All data were processed together following a common pipeline. Specifically, data were trimmed using TrimGalore[23] with default parameters while retaining reads with trimmed length > 25 bases. We then mapped reads from each sample to *Panubis1.0*[25] using STAR 2-pass mapping and a combined splice junction database created from all data during the first round of mapping[74]. Gene-level counts were estimated using HTSeq[75].

Whole blood samples from[14] were sampled, extracted, and sequenced following a different protocol for white blood cell samples collected after 2013. We therefore analyzed the whole blood (n=63) and the white blood cell data sets (n=94) separately. In each case, we filtered for genes with mean TPM ≥ 2 and with counts in at least half of the samples in the dataset, resulting in a set of 11,050 and 11,238 expressed genes in the whole blood and white blood cell data sets, respectively. We then removed genes where local ancestry was nearly fixed (mean anubis ancestry < 10% or > 90%), resulting in 10,615 (whole blood) and 10,798 (white blood cell) genes analyzed for local ancestry effects on gene expression. To assign gene-level local ancestry values to each individual, we calculated the mean number of anubis alleles for a window spanning the gene transcription start site to transcription end site, plus an additional flanking 10 kb on each side. For very short genes, we extended the window size to 50 kb, as local ancestry calls are less reliable for short windows.

Finally, we normalized the gene-level counts matrix for each data set separately using *voom*[76], controlled for library size with *edgeR*[77,78] (version 3.30.3; function *calcNormFactors*), and modeled the normalized gene expression data in *limma*[79] (version 3.44.3; function *lmFit*) as a function of known batch effects (year of sampling and, for white blood cells only, cell type composition summarized using the first two principal components of flow cytometry-based abundance data for $CD3^+CD4^+$ T cells, $CD3^+CD8^+$ T cells, $CD3^-CD20^+$ B cells, $CD3^-CD14^+$ monocytes, and $CD3^-CD16^+$ natural killer cells, as described in Lea et al.[13] and Anderson et al.[15]). We then used the residuals of this model to investigate genetic ancestry effects on gene expression.

*13.2. Testing for additive effects of local genetic ancestry on gene expression levels*

We modeled the normalized, batch-corrected gene expression data using the linear mixed model approach implemented in *emmreml*[80] (version 3.1). We fit fixed effects of local genetic ancestry around each gene, global genetic ancestry, age, and sex, as well as a random effect to control for relatedness between samples. Relatedness estimates (i.e., the K matrix in the mixed model) were calculated from RNA-seq genotype calls (see Anderson et al.[15]) as the covariance of the 0/1/2 genotype matrix. To control for multiple hypothesis testing across genes, we calculated the false discovery rate following[81], against an empirical null distribution derived from 50 permutations of each variable of interest (either global ancestry or local ancestry)[82]. Following the discovery that genome-wide ancestry was not associated with variation in gene expression levels, we further tested whether the effects of genome-wide ancestry were subsumed within the random effect component of the model. However, when we repeated the analysis using a K matrix defined only by pedigree data, which captures familial relatedness but not ancestry, we still did not detect global ancestry effects on gene expression.

*13.3. Testing for non-additive effects of local genetic ancestry on gene expression levels*

Because hybridization has been reported to generate transgressive gene expression patterns in other species pairs (i.e., expression levels outside the range of the parent taxa[83-87]), we also investigated possible non-additive associations between local ancestry and gene expression levels. To do so, we subtracted out the random effect of kinship using *emmreml* before using the *segmented* package[88] (version 1.2-0) to fit a model with different slopes for anubis-like and yellow-like baboons compared to heterozygous individuals. Overall, we found very little evidence for non-additive effects in our sample. Only five genes were both significantly associated with local ancestry and had different slopes for ancestry states 0 to 1 versus 1 to 2 (10% FDR), in either the whole blood or white blood cell data sets. Thus, while local ancestry effects on gene expression in Amboseli are common, most appear to act additively.

We note that our finding of rare non-additive ancestry effects on gene expression may be because anubis baboons and yellow baboons are still relatively closely related (~1.4 million years diverged[8]) compared to the cases reported in the literature[83-87]. Thus, they still share substantial genetic variation and form viable and fertile hybrids. For example, recently diverged zebra finch subspecies (which also hybridize without obvious phenotypic costs) also exhibit gene regulatory divergence but show no evidence of dysregulation in first-generation hybrids[89]. However, it may also be the case that we missed cases of non-additivity that would have been detected in first generation hybrids, as recombination can break apart incompatible combinations in later generations[90].

*13.4. Signatures of selection are stronger near genes with regulatory divergence*

Despite greater genetic divergence between Neanderthals and modern humans, introgressed Neanderthal variants found in modern humans exhibit similar effects on gene expression levels as variants that arose in the modern human lineage[91] (although high frequency Neanderthal alleles may be more likely to affect expression[92]). McCoy and colleagues[91] proposed that the Neanderthal-derived variants that persist in modern humans today represent a non-random subset of those first introduced by admixture, following selection that removed introgressed variants with large effects on gene regulation. However, as less than half the Neanderthal genome is present in modern humans[66,93], it is no longer possible to test this hypothesis across the entire human genome (but see Colbran et al.[94] for predictions in introgression deserts based on variation segregating in modern humans).

It is possible, however, to directly test whether this hypothesis holds for admixture in baboons. If it does, we predict that genes with greater functional divergence (i.e., where local ancestry has large effects on gene expression) will have lower levels of introgressed anubis ancestry than genes that are not functionally diverged. To test this prediction, we rank-ordered all genes in our gene expression analysis by the magnitude of their estimated local ancestry effect (irrespective of sign). Because we generated two estimates for most genes (one for the whole blood RNA-seq data set and one for the white blood cell RNA-seq data set), we first used multivariate adaptive shrinkage (*mashr*[95]) to refine our effect size estimates before calculating the mean effect size across the two data sets. We then calculated the mean ancestry per individual for quantiles of genes with the largest and smallest effect of local ancestry on gene expression (across quantiles from 10-40%; Table S4). We tested whether mean anubis ancestry was different between the two sets, using a paired t-test where mean values for most/least diverged genes were represented for each individual. As reported in the main text and Table S4, we find consistent evidence that functionally diverged genes contained less introgressed ancestry than genes that are not functionally diverged.

We also predict that signatures of selection will be strongest near genes with greater functional divergence. To test this prediction, we used the magnitude of the correlation between recombination rate and introgressed ancestry as a proxy for the strength of selection (as in Fig. 2E, Fig. 3F, "Selection against regulatory divergence"). We first replicated our earlier whole-genome analyses (Section 11.3) for genic regions (transcription start site to transcription end site, plus an additional flanking 10 kb on each side, and extended to a minimum window size of 50 kb). We confirmed that higher local recombination rates also predict higher levels of anubis ancestry in and around protein-coding genes (n=19,050 genes; Spearman's rho = 0.116, p = $1.13 \times 10^{-58}$), with an effect size similar to that estimated for the whole genome (rho = 0.127). This pattern is also detectable among the subset of genes that we tested for ancestry-associated expression (n=10,168; rho = 0.111, p = $1.29 \times 10^{-29}$). In both cases, signatures of selection were again stronger in historically admixed individuals than in recently admixed individuals (all genes: rho = 0.115 vs. rho = -0.013; genes tested for local ancestry-associated gene expression: rho = 0.110 vs. rho = -0.029).

To test whether genes with larger effects of local genetic ancestry on gene expression are associated with stronger signatures of selection against admixture, we calculated the correlation between local recombination rates and anubis ancestry for quantiles of genes with the largest and smallest associations between genetic ancestry and expression (again across quantiles from 10% to 40%). For each quantile comparison, we used a bootstrapping approach to estimate Spearman's rho for the genes with the largest and smallest effect sizes, as well as the difference in rho between these two gene sets (n = 10,000 repetitions). For the more extreme quantiles (≤ 20%), genes with large local ancestry effect sizes consistently exhibited stronger recombination rate-introgressed ancestry correlations than genes with near-zero effect sizes (Fig. 4B; Table S4).

### 14. Predicting anubis ancestry across the genome and through time

#### *14.1. Genomic features predict introgressed ancestry*

To evaluate the capacity to predict the landscape of admixture from genomic features alone, we used *glmnet*[96] (version 4.0-2) to predict introgressed ancestry in 250 kb non-overlapping windows of the baboon genome (n=10,324 windows, after excluding the 50 kb at the chromosome ends and windows where the recombination rate was estimated to be greater than 100x the median recombination rate). Because the predictive accuracy of our models was insensitive to the elastic net mixing parameter, we used a mixing parameter of 0.5 (in *glmnet*, 0 corresponds to ridge regression and 1 corresponds to LASSO[97]). For 200 iterations of *glmnet*, we fit the elastic net model to a random 75% of genomic windows and used the resulting parameter estimates to predict introgressed ancestry in the remaining 25% of windows. We repeated this analysis after permuting introgressed ancestry values to establish a null expectation, and for recent and historical hybrids separately to investigate our relative predictive accuracy.

We included 32 features in our models (Table S5). These features included (i) the number of common genetic variants (minor allele frequency ≥ 5%, based on the full sample of unadmixed reference individuals) and highly differentiated sites ($F_{ST}$ > 0.5-1.0, as described in Section 11.1); (ii) the B statistic (see Section 11.2); and (iii) the local recombination rate based on the anubis baboon genome (see Section 11.3), incorporated as three correlated features (a raw rate, a ranked value, and a $\log_{10}$-transformed value). We also calculated (iv) the proportion of each window that overlapped gene exons, gene bodies, PBMC-predicted enhancers (see Section 11.4), and CpG islands, as well as (v) window GC content. CpG islands were annotated using the *emboss* cpgplot function with default parameters to identify windows of the genome longer than 200 bases with greater than 50% GC content and an observed/expected CpG ratio > 0.6[98,99]. GC content was obtained for each 250 kb window using the nuc function in *bedtools*[65] (v2.25.0). The mean effect size estimate for each feature can be found in Table S5.

*14.2. Genomic features predict the rate of change in anubis allele frequencies*

Anubis ancestry has increased in Amboseli since the early 1980's, when anubis and anubis-like immigrants began arriving in the study population[3] (Fig. 3A). To evaluate whether changes in anubis ancestry since that time are predictable along the genome, we estimated the anubis ancestry proportion in Amboseli for 100 kb genomic windows (excluding the 50 kb at the chromosome ends; n=25,875 total windows), in each chronological year from 1979-2020. We based this calculation on individuals within the Amboseli resequencing data set who were present in the population in that year, according to near-daily demographic records (n=11-228, mean=135.19 resequenced individuals per year). An individual's ancestry was included in the annual population estimate after entering the study population (through birth, immigration, or the onset of observation). They were removed after disappearance from the study population (due to death, dispersal, or because we stopped monitoring their group). For each window, we fit the annual population anubis ancestry as the outcome variable in a linear model with year (1979 – 2020) as a continuous predictor. We estimated the annual change in anubis ancestry ($\Delta_{anubis}$) as the parameter estimate for the year effect.

We then modeled $\Delta_{anubis}$ across windows as a function of the starting anubis ancestry frequency in 1979, mean recombination rate ($\log_{10}$ transformed), weighted $F_{ST}$, and B values (Table S6). Mean recombination rates were calculated as described in Section 11.3, $F_{ST}$ values were calculated as described in Section 7.3, and B values were calculated as described in Section 11.2. We excluded windows in which recombination rate estimates were greater than 100x larger than the median recombination rate and windows that had no variants (so $F_{ST}$ was not calculated), resulting in a final data set of 25,797 windows. We also confirmed that estimates of model effects were generally stable across window sizes from 35 - 250 kb.

### Supplementary Figures

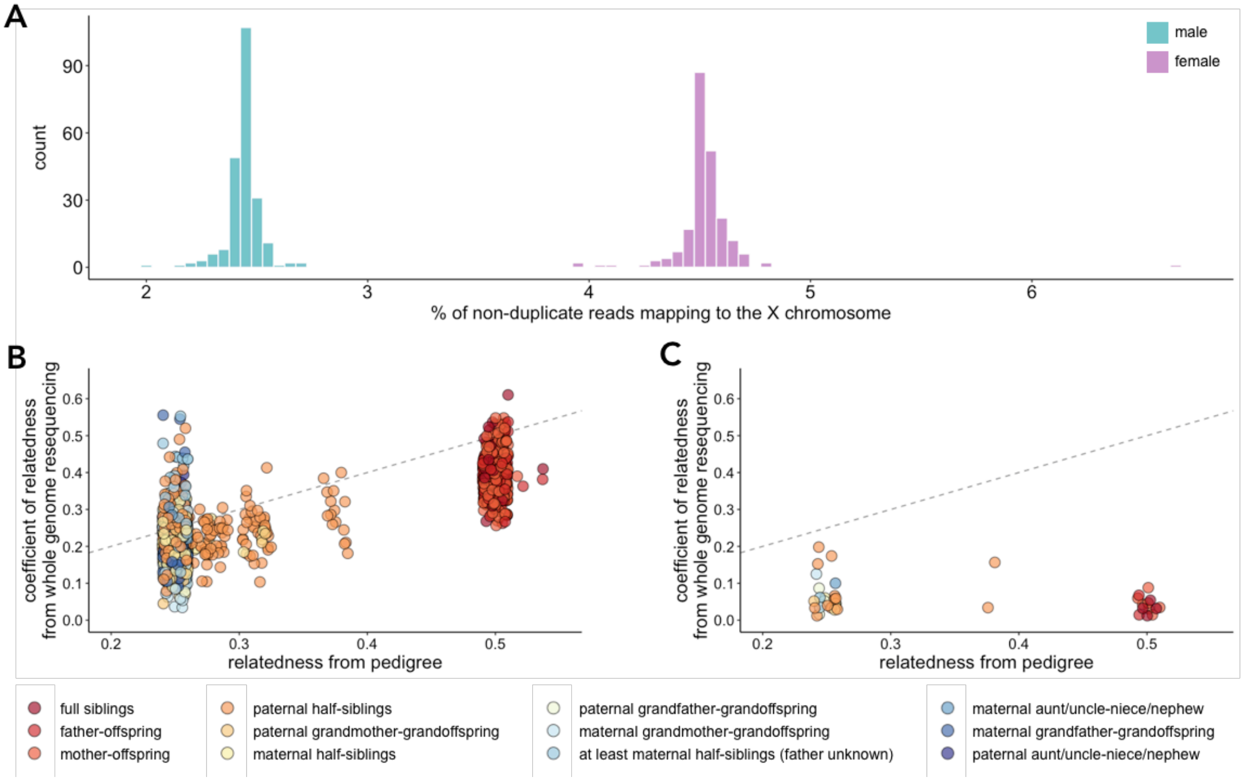

**Figure S1. Quality control for individual identity in the Amboseli resequencing data. (A)** For sex, we confirmed that resequencing data from females (pink) had approximately 2x the percentage of non-duplicate mapped reads assigned to the X chromosome relative to males (blue). **(B-C)** Relationship between relatedness estimates from the Amboseli Baboon Research Project pedigree and from genome resequencing using *lcMLkin*[28] for pairs of individuals with expected pedigree-based relatedness ≥ 0.25 (n=382 unique individuals). **B** shows dyads where both individuals were correctly identified (n=376 unique individuals), resulting in a strong correlation between pedigree and sequencing-based relatedness (Pearson's r = 0.789, $p < 10^{-300}$). **C** shows dyads which include at least one of 6 incorrectly identified individuals, where pedigree and sequencing-based relatedness estimates are not correlated (Pearson's r = -0.244, p = 0.129). Cases of expected pedigree-based relatedness equal to 0.5 (e.g., parent-offspring, full siblings) are colored in dark orange or red. The grey dashed line corresponds to the y = x line. The considerable noise in this relationship likely reflects noise in relatedness estimates from low coverage resequencing data, structure from the admixed nature of the population, and some degree of cryptic relatedness not apparent in the pedigree data.

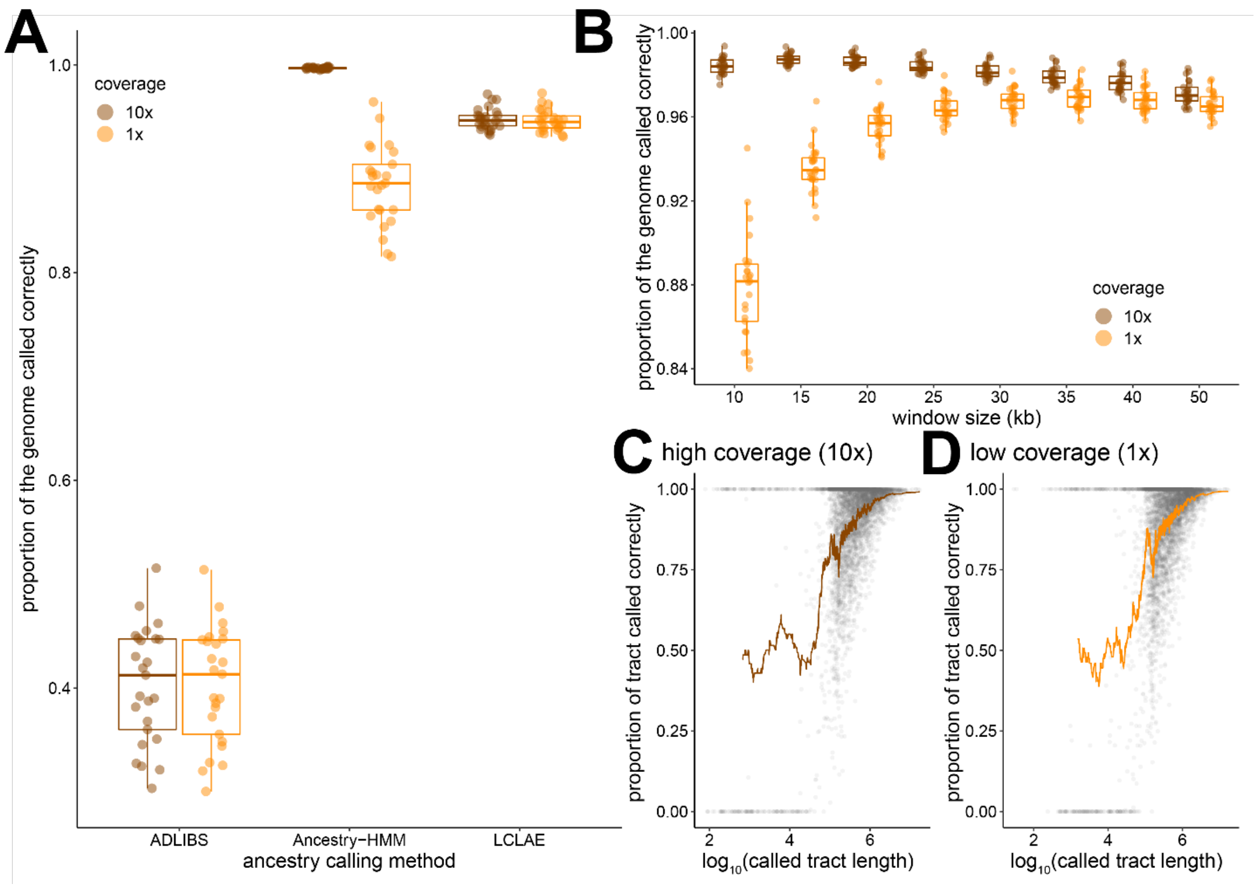

**Figure S2. Accuracy of local ancestry assignment. (A)** Comparison of local ancestry calling using AD-LIBS[36], Ancestry HMM[35], and LCLAE[6] with both high (10x, brown) and low (1x, orange) coverage data for 25 simulated individuals. **(B)** The proportion of the genome called correctly using LCLAE as a function of the window size (for calling ancestry) and sequencing coverage. The smallest window sizes are less accurate using low coverage data because few ancestry informative sites are available for making assignments, while the largest window sizes are less accurate for both levels of coverage because they miss small ancestry tracts. **(C-D)** The relationship between called tract length and accuracy in calling local ancestry with LCLAE for high (C) and low (D) coverage data. Each point represents an ancestry tract called for a simulated individual, and the brown (C) and gold (D) lines represent the moving mean accuracy for non-overlapping sets of 70 ancestry tracts. Tracts less than 1 kb perform little better than chance. In contrast, longer tracts are close to 95% accurate, with the main source of error due to missed small ancestry tracts that are contained within large tracts.

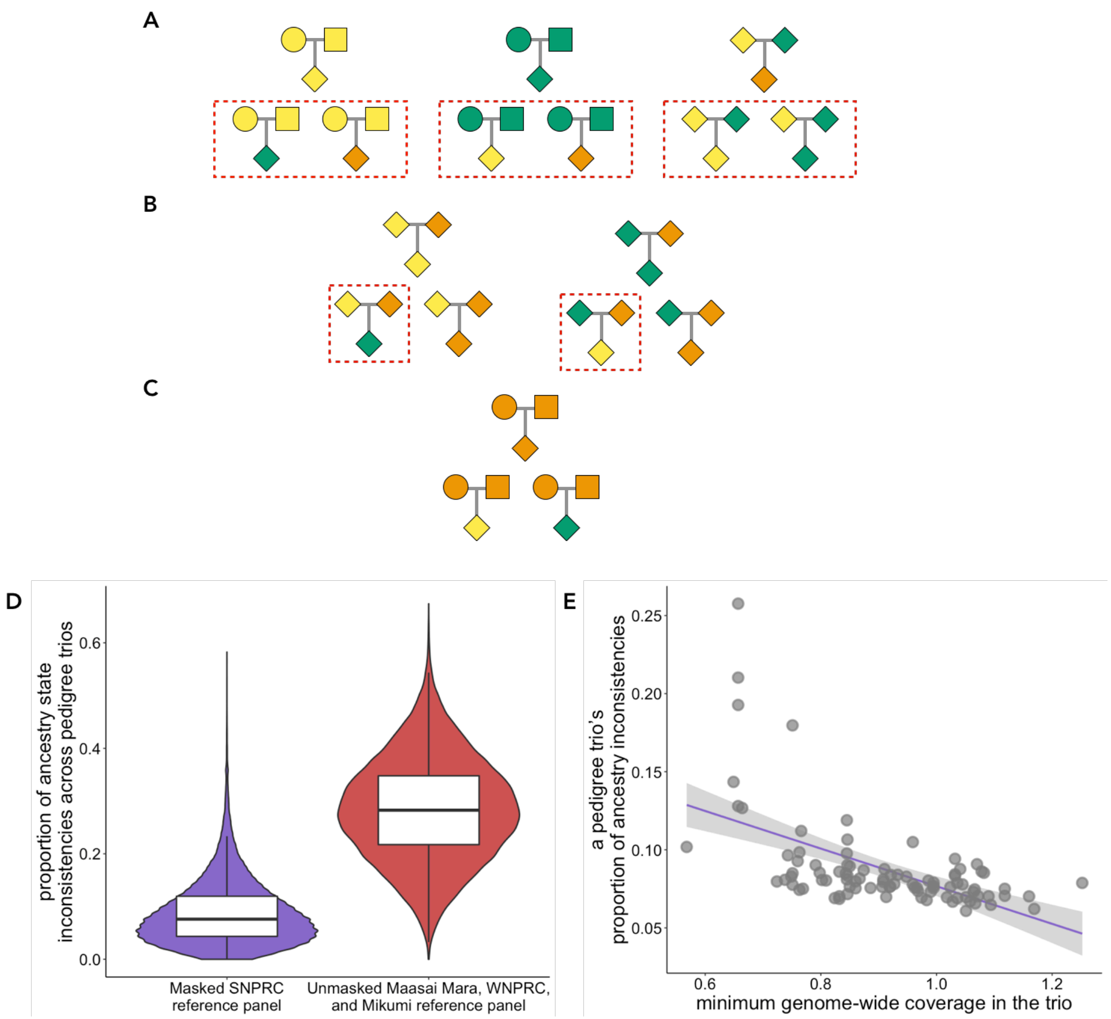

**Figure S3. Local ancestry call consistency within pedigree trios. (A-C)** Ancestry configurations for pedigree trios. Pedigree trios with internally inconsistent calls are shown inside red dashed boxes. Females = circles, squares = males, diamonds = males or females (i.e., the sex of the individual does not matter); yellow = homozygous yellow baboon ancestry, orange = heterozygous yellow-anubis baboon ancestry, green = homozygous anubis baboon ancestry. **(A)** Pedigree trios in which both parents are homozygotes. **(B)** Pedigree trios in which one parent has homozygous ancestry and the other parent has heterozygous ancestry. **(C)** Pedigree trios in which both parents are heterozygotes. **(D)** Proportion of ancestry state inconsistencies per site across pedigree trios using different reference populations for yellow baboon and anubis baboon ancestry (the SNPRC reference panels are masked for introgressed segments; see Supplementary Methods Section 6.2). Note that the SNPRC reference panel consists of only medium to high-coverage sequences, whereas the Maasai Mara/WNPRC/Mikumi reference panel contains a mix of low and high coverage sequence data. **(E)** The proportion of sites with internally inconsistent calls, within trios, is strongly negatively correlated with the minimum genome-wide coverage among members of the trio (Pearson's $r = -0.553$, $p = 1.07 \times 10^{-8}$).

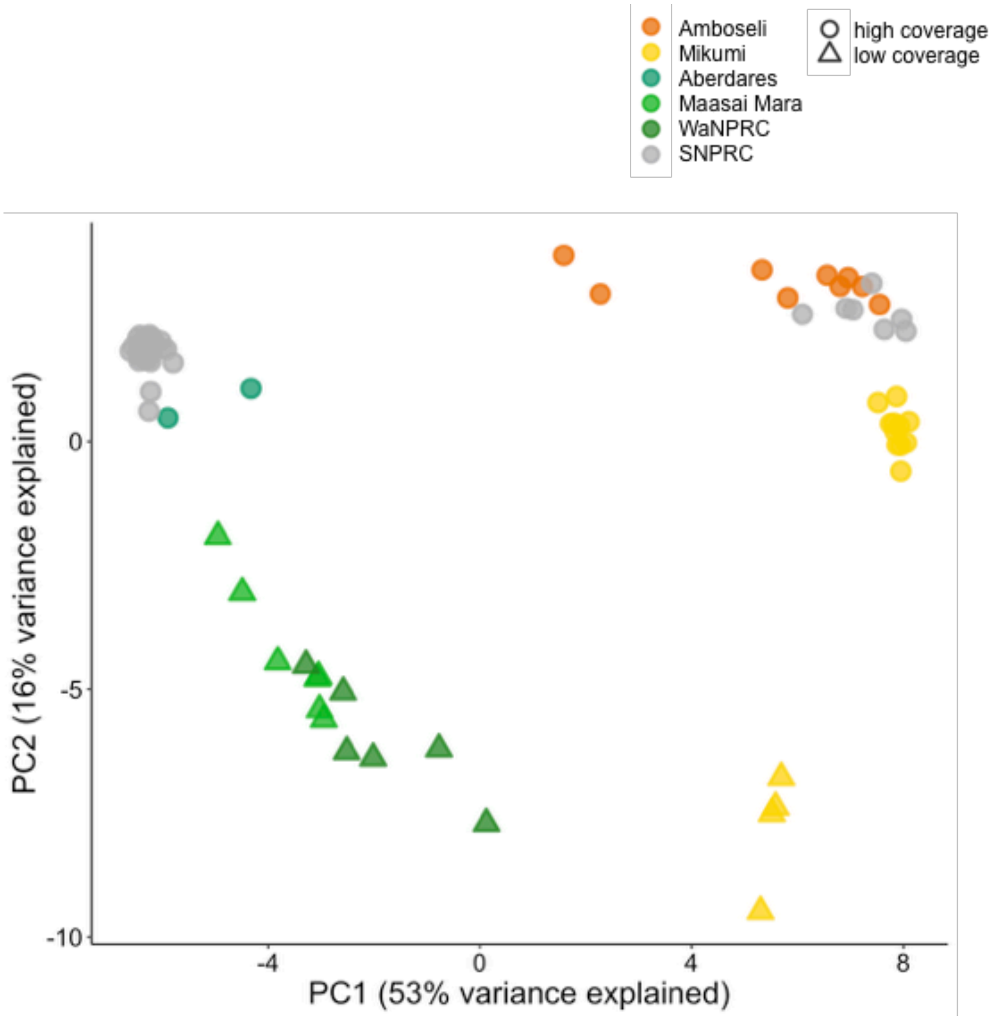

**Figure S4. The structure of genetic variation between yellow, anubis, and hybrid baboons.** Principal components analysis of genotypes from high coverage samples (SNPRC: grey circles; the Aberdares region: green circles; Mikumi: yellow circles; Amboseli: orange circles) and low coverage samples (Mikumi: yellow triangles; Maasai Mara: light green triangles; WNPRC: dark green triangles). Ancestry differences predominate on PC1, but sequencing coverage is also correlated with PC2.

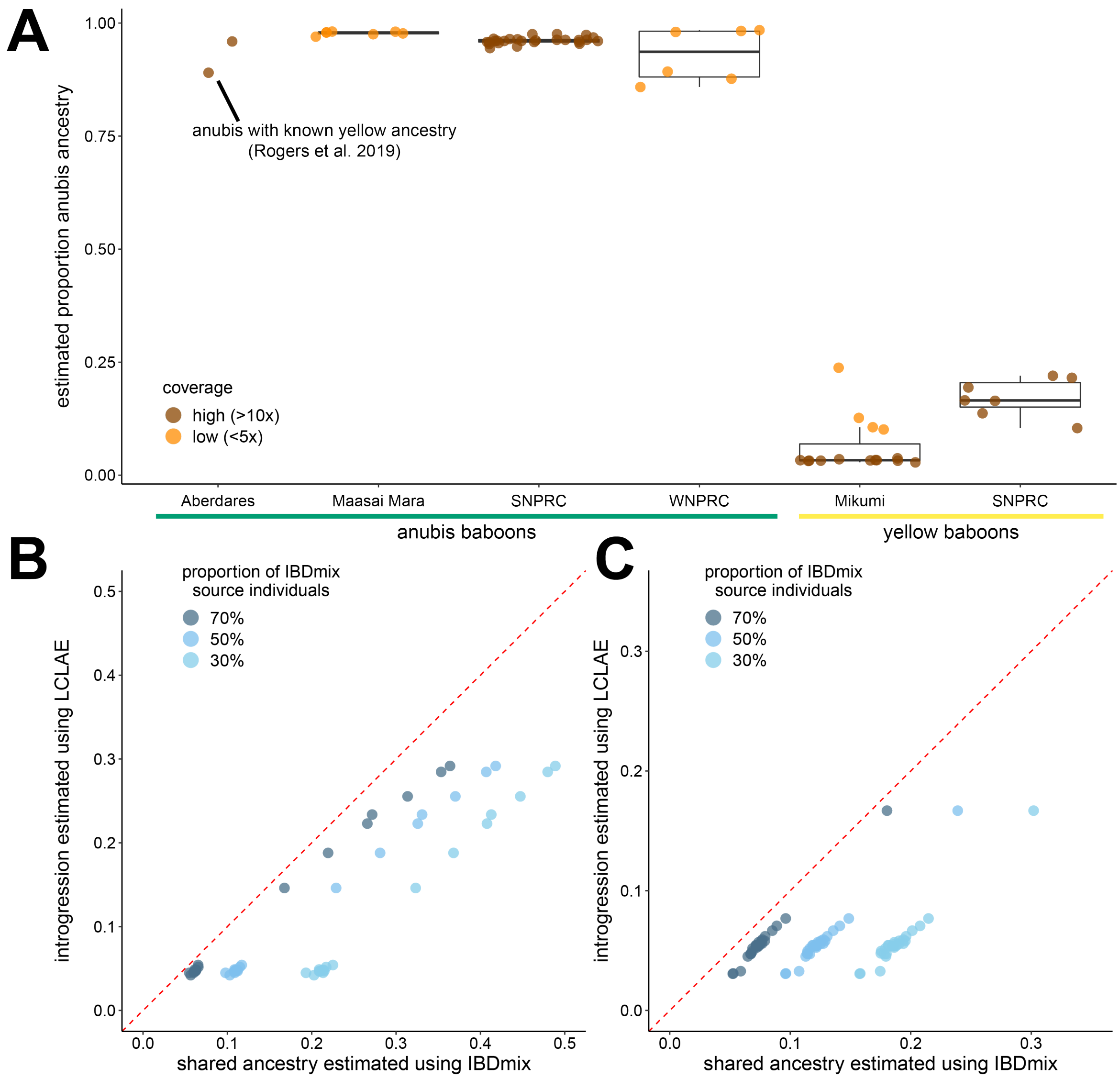

**Figure S5. Admixture in yellow and anubis baboons outside of Amboseli. (A)** Genome-wide proportion of anubis ancestry inferred per baboon using LCLAE. Estimates for high coverage individuals are shown in brown, while low coverage individuals are shown in orange. **(B-C)** Estimates of local ancestry are strongly correlated between IBDmix and LCLAE. **B** shows the proportion of anubis ancestry identified in yellow baboons; **C** shows the proportion of yellow ancestry identified in anubis baboons. Darker points represent the more conservative version of IBDmix results, requiring that a region of the genome be identified as IBD using 70% of potential source individuals. Successive lighter points represent increasingly liberal standards (50% or 30% of source individual agreement). Dashed red lines correspond to the y = x line, showing that IBDmix tends to overestimate shared ancestry relative to LCLAE.

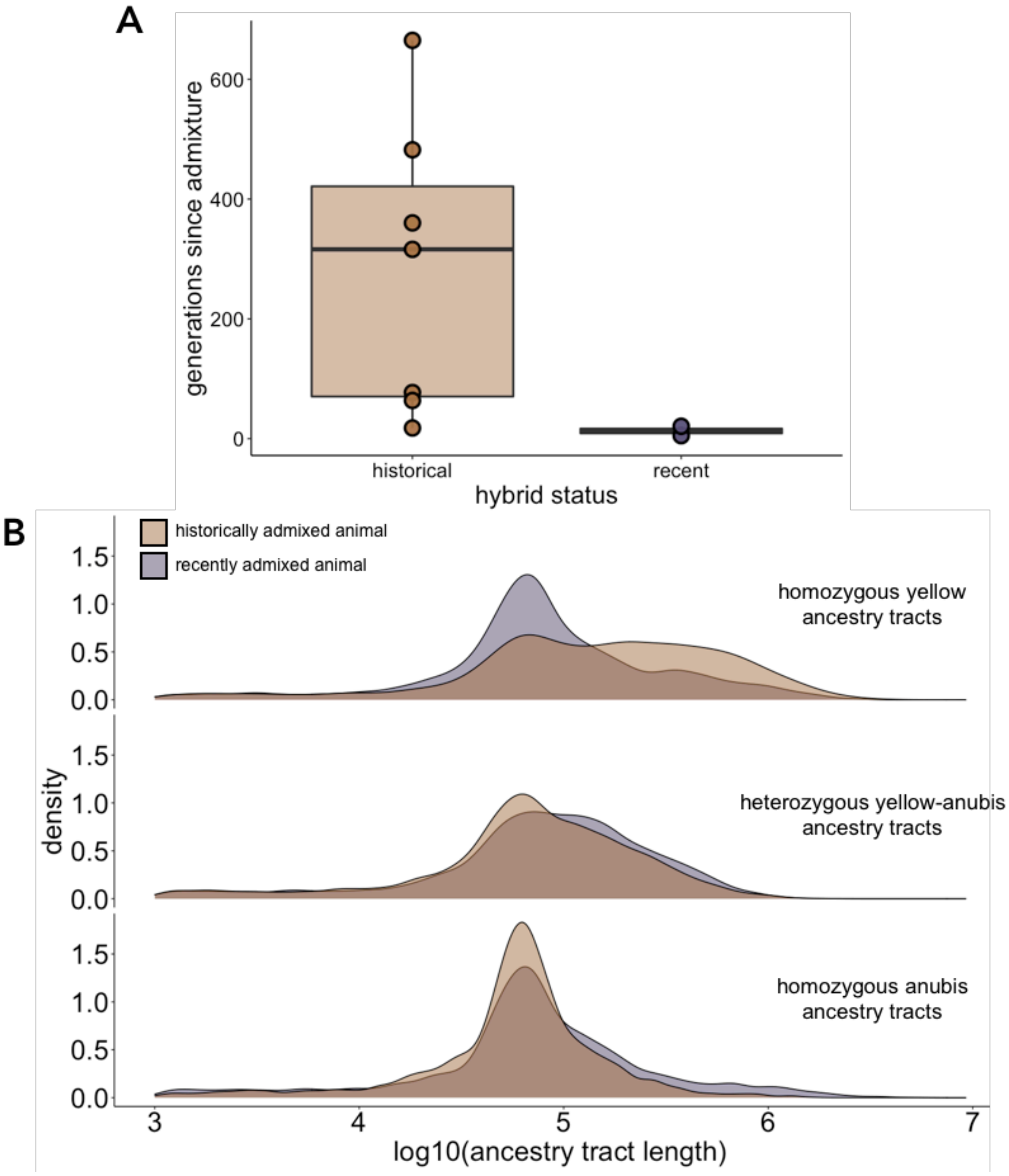

**Figure S6. Estimated timing of admixture in Amboseli baboons. (A)** *DATES*[49] estimates for the timing of admixture for Amboseli individuals sequenced at high coverage. Individuals were assigned to “recent” or “historical” hybrid categories based on whether they had a known anubis or anubis-like ancestor (Supplementary Methods Section 1.1). **(B)** The distribution of ancestry tract lengths for a recently admixed animal in (A) with an estimate of 4.99 generations since admixture (purple colored distributions) versus a historically admixed animal in (A) with an estimate of 316.12 generations since admixture (brown colored distributions).

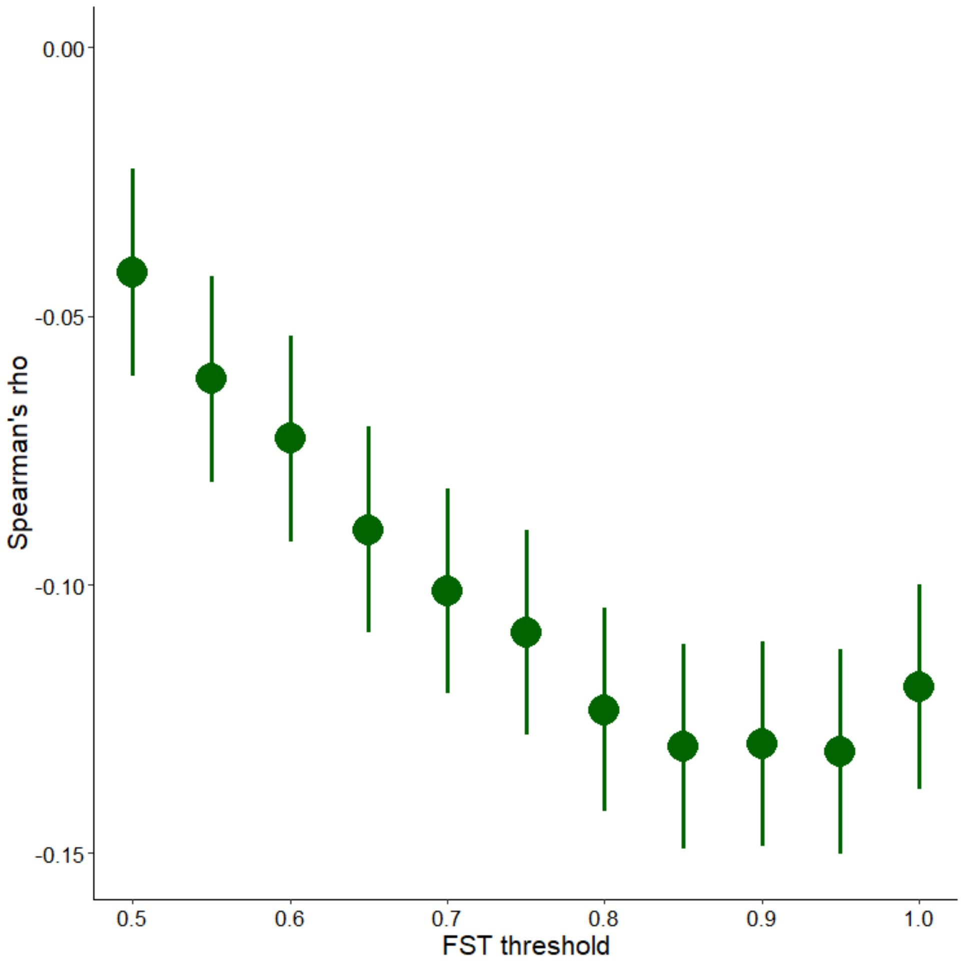

**Figure S7. Introgressed ancestry is reduced in highly diverged regions of the genome.** Spearman's rho for the correlation between introgressed ancestry and the number of highly differentiated sites in a 250 kb region of the genome, across variable minimum $F_{ST}$ thresholds (x-axis) for defining highly differentiated sites. Error bars show the standard error of rho. For all threshold values of $F_{ST}$, rho is significant at $p < 2.5 \times 10^{-5}$.
